# Brieflow: An Integrated Computational Pipeline for High-Throughput Analysis of Optical Pooled Screening Data

**DOI:** 10.1101/2025.05.26.656231

**Authors:** Matteo Di Bernardo, Roshan S. Kern, Ana Karla Cepeda Diaz, Alexa Mallar, Samuel J. Choi, Andrew Nutter-Upham, Sebastian Lourido, Paul C. Blainey, Iain Cheeseman

## Abstract

Optical pooled screening (OPS) has emerged as a powerful technique for functional genomics, enabling researchers to link genetic perturbations with complex cellular morphological phenotypes at unprecedented scale. However, OPS data analysis presents challenges due to massive datasets, complex multi-modal integration requirements, and the absence of standardized frameworks. Here, we present Brieflow, a computational pipeline for end-to-end analysis of fixed-cell optical pooled screening data. We demonstrate Brieflow’s capabilities through reanalysis of a CRISPR-Cas9 screen encompassing 5,072 fitness-conferring genes, processing more than 70 million cells with multiple phenotypic markers. To accelerate biological interpretation, we additionally present MozzareLLM, a framework leveraging large language models to identify biological processes within phenotypic clusters and prioritize gene candidates for experimental validation. Our combined analysis recovered coherent biological modules missed by existing analytical approaches, including five core mitochondrial sub-programs that were absent from the original study. The modular design and open-source implementation of Brieflow facilitates the integration of novel analytical components while ensuring computational reproducibility and improved performance for the use of high-content phenotypic screening in biological discovery.

## Background

Functional genomics has been transformed by high-throughput screening technologies that enable the systematic interrogation of gene function. Optical Pooled Screening (OPS) has emerged as a powerful technique for elucidating gene-phenotype relationships by combining pooled genetic perturbations with high-resolution cellular imaging^1^. In a typical OPS workflow, cells are transduced with a pooled library of genetic perturbations (e.g., CRISPR-Cas9), each identified by a unique barcode sequence (Fig. 1a, panel I). Following selection and fixation, cells undergo high-content imaging to capture phenotypic features followed by *in situ* sequencing to identify barcode sequences (Fig. 1a, panel II).

**Fig. 1.**
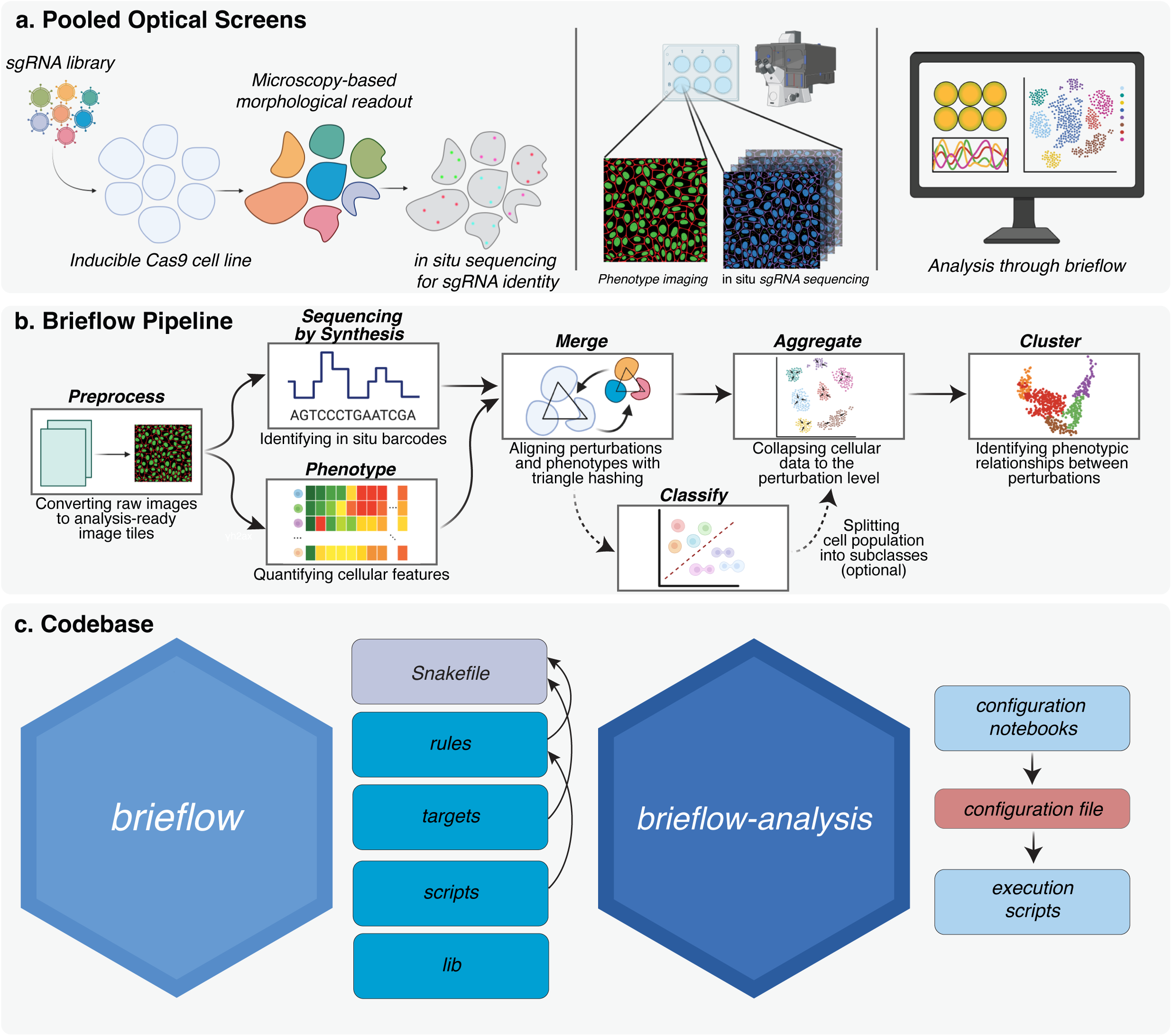
Brieflow: An integrated computational framework for optical pooled screening analysis. **a.** Schematic of the optical pooled screening workflow. Cells expressing inducible Cas9 are transduced with an sgRNA library, perturbed, and subjected to both phenotypic imaging and in situ sgRNA sequencing, followed by computational analysis using Brieflow. **b.** The Brieflow pipeline architecture comprising seven integrated modules: Preprocess converts raw microscopy files to standardized formats; Sequencing-by-Synthesis identifies guide barcodes; Phenotype extracts cellular features; Merge integrates genotype and phenotype data using triangulation-based spatial registration; Classify optionally partitions cells into relevant subpopulations; Aggregate condenses single-cell data to perturbation-level statistics; and Cluster identifies functional relationships between perturbations. **c.** Implementation architecture. Left: the Brieflow codebase with modular structure, where a high-level Snakefile imports rules and targets from each module, rules execute individual Python scripts, and scripts call functions defined in the library. Right: the complementary Brieflow-analysis template, where a configuration notebook allows users to test and set parameters that are automatically written into the configuration file, which is then employed by execution scripts that call the Snakefile with appropriate module-specific parameters.

Unlike traditional pooled screening approaches that rely on simple readouts such as cell viability, OPS captures rich, multidimensional phenotypic data while maintaining advantages including increased throughput and reduced reagent consumption. Furthermore, OPS integrates seamlessly into the cell biologist’s toolkit, effectively scaling standard imaging-based analyses to enable systematic investigation of stains, antibodies, and fluorescent reporters at unprecedented throughput. OPS has been successfully deployed for diverse applications, including the discovery of novel antiviral response pathway components^2^, the identification of cellular factors regulating transcription factor dynamics^3^, and the establishment of previously unknown connections between genes and complex morphological phenotypes^4^.

The scale and complexity of OPS data present formidable computational obstacles. These terabyte-size datasets often contain information from multi-channel fluorescence images of millions of individual cells, each characterized by thousands of phenotypic parameters. The integration of *in situ* sequencing and phenotypic measurements necessitates accurate matching of barcode identities with corresponding phenotypic features across different imaging modalities with distinct optical characteristics and resolution limitations. Normalization across experimental batches and conditions is essential to minimize technical variation and enable reliable biological comparisons. The implementation of OPS has also been hampered by a lack of standardized, comprehensive codebases. Although powerful computational methods have been developed that leverage processed OPS data for tasks such as perturbation prediction and phenotypic modeling^5,6,7^, the field lacks both a standardized analysis pipeline accessible to experimental biologists and a consistent output format for training computational models. The processing pipelines necessary to generate these inputs—from barcode demultiplexing and cell-perturbation assignment through phenotype feature extraction—have remained fragmented, lab-specific, and often require manual intervention at multiple stages, slowing adoption by wet-lab researchers and limiting the standardized datasets available for downstream modeling efforts^8^. Finally, the immense size of raw image datasets makes conventional data sharing impractical and incomplete documentation of analysis parameters undermines reproducibility.

To address these challenges, we developed Brieflow, a computational pipeline designed specifically for end-to-end analysis of fixed-cell OPS datasets. Brieflow provides a standardized framework for processing and analyzing optical pooled screening data, offering a critical pathway to incorporate complex cellular morphologies into emerging in silico cell models. To demonstrate the improvements offered by our approach, we benchmarked Brieflow through a comprehensive reanalysis of a large-scale CRISPR-Cas9 screen comprising over 5,000 fitness-conferring genes in HeLa cells^9^, illustrating its efficacy in processing and extracting biological insights from terabyte-scale OPS datasets. To enable the advanced interpretation of these datasets, we introduce MozzareLLM, a specialized large language model framework for biological interpretation that automatically analyzes phenotypic clusters for their functional processes and identifies promising gene candidates for experimental validation. By enabling reproducible, consistent analysis across diverse experimental contexts with improved quality and precision, this pipeline represents an essential step toward integrating the morphological dimension of cellular function into comprehensive computational models of cell biology.

## Results

### Brieflow: An Integrated Framework for OPS Data Analysis

The analysis of OPS datasets is a multi-step process that requires specialized computational tools to extract, integrate, and interpret the multidimensional phenotypic data. To enable the analysis of these data, we developed a computational pipeline that we term Brieflow (Fig. 1a, panel III). Brieflow provides a unified framework that processes data from raw images to generate biological insights. Brieflow employs a modular architecture comprising seven integrated components (Fig. 1b): Preprocess, Sequencing-by-Synthesis, Phenotype, Merge, Classify, Aggregate, and Cluster. As described in detail below, the Preprocess module converts raw images to analysis-ready image tiles; Sequencing-by-Synthesis identifies *in situ* barcodes of genetic perturbations; Phenotype quantifies cellular features from high-content imaging; Merge aligns perturbations and phenotypes using triangle hashing; Classify partitions cells into relevant subpopulations; Aggregate collapses cellular data to the perturbation level; and Cluster identifies phenotypic relationships between perturbations.

Brieflow’s modular architecture enables flexible workflows through independent components with well-defined inputs and outputs, allowing execution of either the complete pipeline or individual modules as needed. The implementation leverages Snakemake-based workflow management, which automatically determines processing steps based on file dependencies for incremental updates (Extended Data Fig. 1), distributes independent tasks across available computational resources, and captures complete analytical provenance in a version-controlled format for reproducibility^10^. The architecture comprises library code, Python scripts, Snakemake rules, and targets according to a standardized project development template (Fig. 1c, panel I). The Brieflow-analysis template repository provides a hierarchical, YAML-based configuration system that separates experimental parameters from processing logic, with configuration Jupyter notebooks that guide users through parameter selection for each module and execution scripts optimized for both local and high-performance computing environments (Fig. 1c, panel II). As a case study, we use Brieflow to reanalyze the OPS data from Funk et al. 2022 with the Cas9-based targeting of ∼5000 fitness conferring genes in human HeLa cells, herein designated as the ‘Vesuvius’ dataset, with specific findings described for each step below.

### Preprocess: Conversion of raw microscopy files to standardized tiled images

The analysis of complex OPS datasets first requires standardization of diverse microscopy data formats into a unified, analysis-ready structure. The Preprocess module addresses initial challenges of OPS data analysis by providing tools for data conversion, metadata extraction, and the generation of appropriate illumination correction functions. The module supports multiple input formats through a configurable interface: users specify their file type (ND2 or TIFF) and associated metadata structure, enabling compatibility with diverse microscopy systems including Nikon (ND2), Phenix (TIFF), and Squid (TIFF) platforms. For TIFF files, users provide accompanying metadata files. The module also supports multiple-round phenotyping via iterative staining, with each round treated as an independent phenotyping cycle that is processed separately before downstream integration. We note that the current implementation assumes one-to-one tile correspondence within phenotyping rounds and sequencing-by-synthesis cycles, which is satisfied by most standard microscope configurations.

Illumination correction is applied to normalize signal intensity across the field of view (Extended Data Fig. 2)^11^. In our reanalysis of the ‘Vesuvius’ screen, the Preprocess Module processed 2,049 ND2 input files, generating 168,165 SBS tiles (across 11 sequencing rounds) and 58,926 phenotype tiles. The Preprocess module establishes a flexible foundation for downstream analysis by transforming heterogeneous microscopy data into standardized, efficiently-structured image tiles.

### Sequencing-by-synthesis: Accurate identification of genetic perturbations

Linking cellular phenotypes to their causative genetic perturbations requires identification of barcodes within cells from *in situ* sequencing images. The Sequencing-by-synthesis module (Fig. 2a) processes multi-cycle sequencing images through alignment, spot detection, base calling, cell segmentation, and barcode-to-cell mapping, generating detailed quality metrics including read mapping rates and cell assignment statistics at each stage (Fig. 2b).

**Fig. 2.**
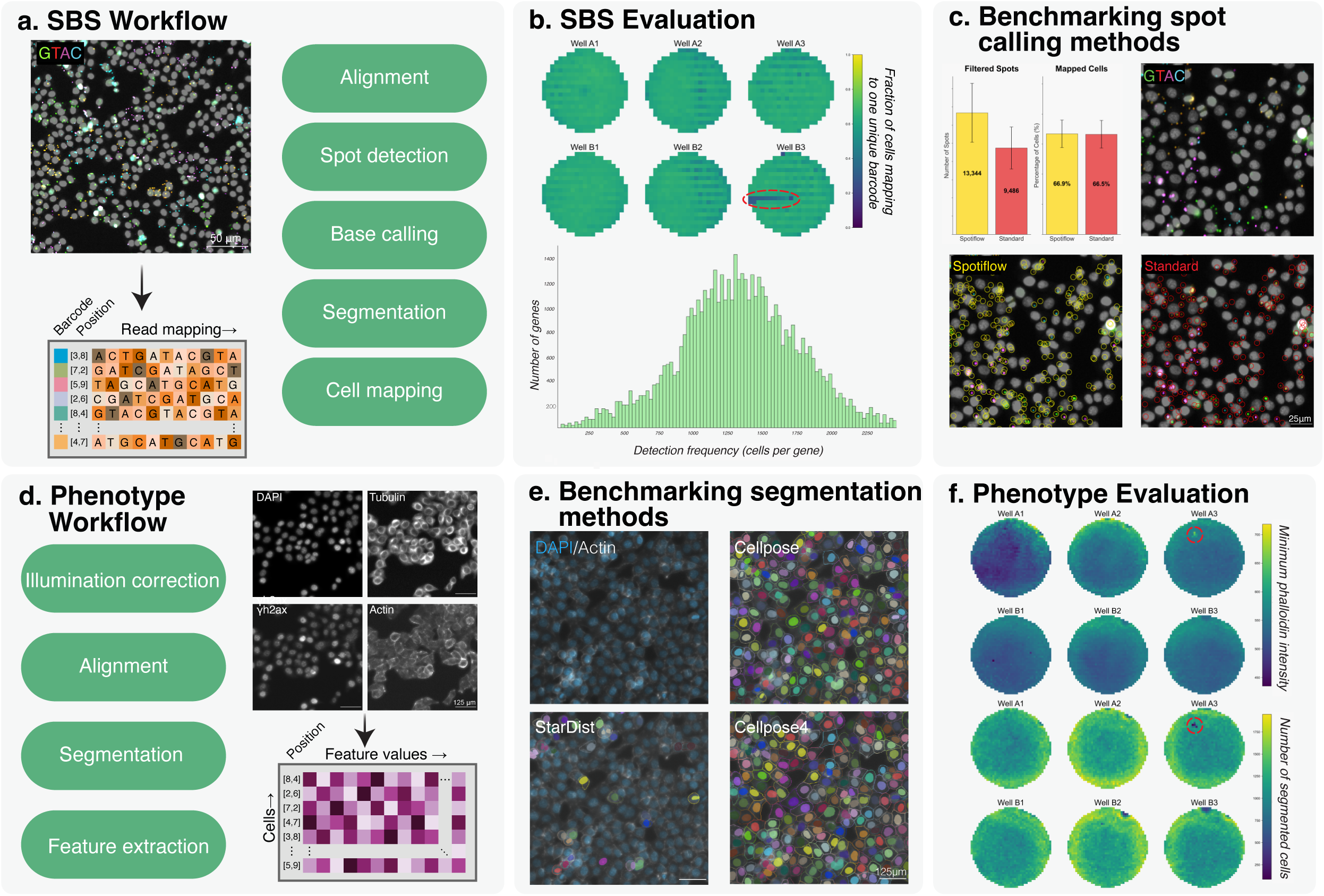
Sequencing-by-Synthesis and Phenotype modules for analysis of optical pooled screening data. **a.** Workflow of the Sequencing-by-Synthesis module. In situ sequencing images undergo multi-cycle alignment, spot detection, base calling, cell segmentation, and barcode-to-cell mapping, generating a comprehensive output table with cellular barcode assignments and per-base and per-barcode mapping statistics. **b.** Quality assessment metrics for plate 1. Top: heatmap showing the fraction of cells mapping to a single gene, enabling identification of areas with low spot signal (circled). Bottom: distribution of cells across genetic perturbations. **c.** Comparison of spot detection methods on a representative tile. Spotiflow detected more spots per tile than the standard method with a threshold of 400 (13,334 vs. 9,486) but resulted in comparable gene mapping efficiency (66.9% vs. 66.5%). **d.** Phenotype module workflow. Multi-channel fluorescence images undergo illumination correction, channel alignment, segmentation, and feature extraction, quantifying morphological and functional characteristics across cellular compartments. **e.** Benchmarking of segmentation approaches on a representative tile. Cellpose achieved superior segmentation of HeLa cells by qualitative inspection. **f.** Quality assessment of phenotypic measurements for plate 1. Top: minimum phalloidin intensity used for cytoplasmic segmentation quality assessment. Bottom: cell count distribution, demonstrating correlation between staining anomalies and decreased segmentation efficiency (circled).

The module offers two distinct approaches for identifying sequencing spots: a standard method based on classical image processing signal detection and a deep learning-based method using Spotiflow^12^. In benchmarking analysis across eight randomly selected tiles (one per plate), the standard method detected substantially more initial peaks than Spotiflow (48,133 vs. 13,344 peaks on average per tile), though after quality filtering, the standard method yielded 9,486 high-quality reads compared to 13,344 per tile for Spotiflow (Fig. 2c). Critically, read mapping fractions were higher with the standard method (71.8% vs. 57.6%), although the proportion of cells successfully assigned to a single gene target remained comparable between approaches (66.5% vs. 66.9%). Processing with Spotiflow required substantially more time, averaging 40.3 seconds per tile compared to just 0.87 seconds with the standard method (Extended Data Fig. 3a). Given the superior mapping rates and dramatically faster processing speed and lower memory usage (Extended Data Fig. 3b), we employed the standard method for our reanalysis of the Vesuvius screen.

Following spot detection, base calling corrects for channel cross-talk using configurable normalization strategies. For our analysis, we selected median-based normalization on the basis of its superior correction of A-C crosstalk (Extended Data Fig. 3c). Reads are mapped to a reference barcode library with support for error-tolerant matching and combinatorial barcode schemes used in PerturbView and Zombie *in situ* sequencing approaches^13^. A grid search utility evaluates combinations of detection parameters on representative tiles, reporting spot count, mapping rate, and cell assignment rate for each combination, enabling users to identify optimal settings before full-scale processing. In our reanalysis of the Vesuvius dataset, we identified a total of 74,406,684 cells with 79.2% (58,980,065) containing a sgRNA barcode, and 57.0% (42,371,543) successfully assigned to a unique gene target.

To validate Brieflow’s generalizability, we applied the SBS module to an independent dataset generated using three known barcode constructs sequenced across three cycles using T7-based chemistry. Brieflow accurately mapped 78.2% of the segmented cells uniquely to a single construct (Extended Data Fig. 3d). PerturbView-style *in situ* sequencing produces single high-intensity nuclear peaks^13^ rather than distributed cytoplasmic signal; both characteristics are accommodated through Brieflow’s configurable parameters for peak detection prioritization and compartment-specific read assignment. Together, the Sequencing-by-synthesis module transforms complex multi-cycle imaging data into high-confidence genetic perturbation assignments for each cell.

### Phenotype: Comprehensive cellular feature extraction

Capturing the phenotypic impact of genetic perturbations requires the extraction of comprehensive multi-dimensional cellular features from high-content imaging data. To achieve this, the Phenotype module (Fig. 2d) implements a systematic workflow beginning with specimen-specific illumination correction to normalize signal across the field of view, followed by alignment of fluorescence channels to correct for chromatic aberrations, cellular segmentation to identify nuclear and cellular boundaries, and finally feature extraction to quantify morphological and functional characteristics across multiple cellular compartments.

Channel alignment is performed using phase cross-correlation for sub-pixel precision, with integrated notebook visualizations enabling manual inspection of registration accuracy. For high-magnification or multi-round imaging contexts where automated alignment may be insufficient, the module supports a two-stage approach combining user-specified coarse offsets with automated sub-pixel refinement. Custom channel-specific offsets can additionally be specified to correct for systematic chromatic aberrations.

For cellular segmentation, Brieflow supports multiple deep learning-based approaches: Cellpose3^14^ (cyto3 model), Cellpose4^15^ (CPSAM model, integrating the Segment Anything foundation model), and StarDist^16^. Segmentation masks are applied across all channels for feature extraction, with no requirement to segment each channel individually. Notebook visualizations based on visual inspection and the proportion of segmented nuclei corresponding to segmented cells enable users to test segmentation approaches on representative tiles before committing to full-scale processing. In comparative benchmarking on one random imaging tile per plate (8 tiles total), Cellpose (cyto3) achieved high retention rates (93.0% nuclei, 93.7% cell retention) with moderate processing time (186 seconds per tile). Cellpose4 (CPSAM) achieved comparable segmentation but with expectedly longer processing times (1,161 seconds per tile). However, CPSAM offers strong performance for non-traditional morphologies such as elongated RPE1 cells, making it a valuable alternative for challenging cell types. StarDist was most computationally efficient (110 seconds per tile) but achieved lower retention rates (16.7% nuclei, 35.1% cell retention), with many detected objects lost during downstream reconciliation (Fig. 2e, Extended Data Fig. 4a-d). Based on these results, we selected Cellpose (cyto3) for the re-analysis of the Vesuvius screen.

For feature extraction, Brieflow supports two configurable approaches: a CellProfiler-inspired^17^ pure Python emulator developed to address integration challenges between Java-based CellProfiler and Brieflow’s Snakemake workflow, and a package (cp_measure) developed by the CellProfiler team^18^. The emulator extracts four feature categories — intensity, texture, shape, and correlation — across nucleus, cell, and cytoplasm compartments; a comprehensive feature dictionary is provided in the Brieflow repository. Benchmarking against cp_measure revealed strong agreement for interpretable intensity and shape features, although features sensitive to local variations — particularly radial distribution features — showed divergence between implementations (Extended Data Fig. 4g-h). These discrepancies likely reflect differences in implementation details for complex multi-step computations rather than fundamental algorithmic differences. Furthermore, the cp_measure implementation required an average processing time of close to 5 hours per tile, which is impractical for high-content screens. In our reanalysis, the Phenotype module segmented 71,629,095 cells with 1,651 morphological measurements extracted per cell (Fig. 2f). Together, the Phenotype module converts raw fluorescence images into rich quantitative profiles that capture the morphological consequences of genetic manipulations, informing the functional contribution of each corresponding gene product.

### Merge: Integration of genetic and phenotypic data

Connecting a genetic perturbation with its phenotypic effect necessitates spatial registration between different imaging magnifications captured during the optical pooled screening workflow. The Merge module integrates data from the Sequencing-by-synthesis and Phenotype modules, creating a unified dataset that links genetic perturbations with their corresponding phenotypic effects (Fig. 3a). Because OPS workflows typically image barcodes and phenotypes at different magnifications and potentially even on different microscopes, direct pixel-to-pixel correspondence cannot be assumed. Instead, the merge process leverages the spatial pattern of cells themselves as fiducial markers: cells occupy consistent physical positions regardless of imaging magnification, so the geometric relationships between neighboring cells can be used to identify corresponding regions across modalities. The implementation uses Delaunay triangulation to create hash-based descriptors of local cell arrangements that can be matched across imaging modalities despite differences in scale and orientation.

**Fig. 3.**
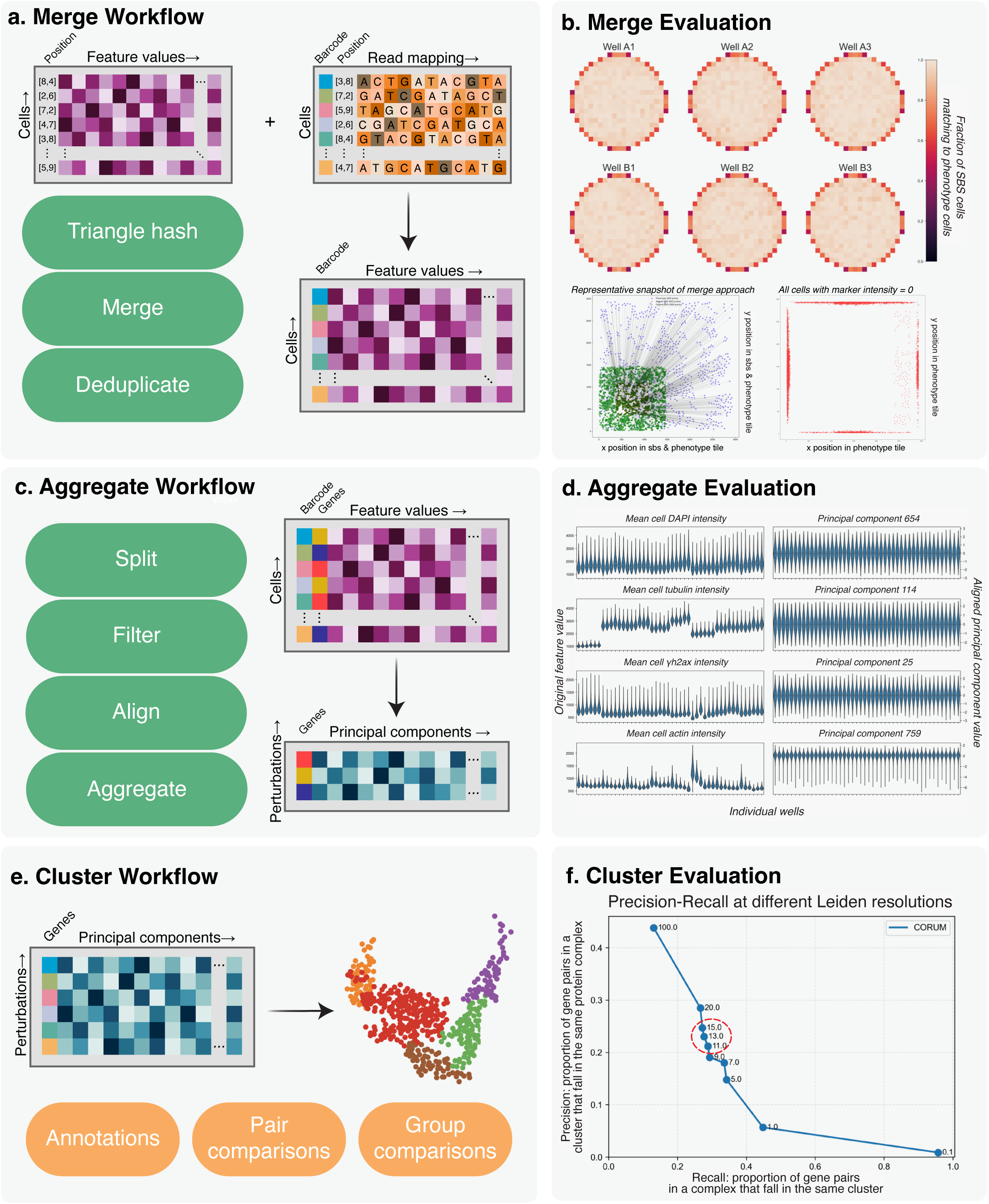
Merge, Aggregate, and Cluster modules for integrating genetic and phenotypic data. **a.** Merge workflow integrating phenotypic features with genetic perturbation data. Phenotypic measurements (left) and barcode mapping statistics (right) are combined through triangle hashing for spatial registration, cell merging, and deduplication to resolve multiple matches, producing a unified dataset (bottom) linking genetic perturbations with phenotypic signatures. **b.** Quality assessment of Merge performance for plate 3. Top: heatmaps showing the fraction of SBS cells matching to phenotype cells across all wells. Bottom left: representative snapshot of spatial registration between SBS and phenotype cell positions for a single well, with axes corresponding to tile coordinates. Bottom right: spatial distribution of cells with minimum marker intensity of zero, which are deprioritized during deduplication; these predominantly occur at tile boundaries due to optical edge effects. **c.** Aggregate workflow for transforming single-cell data into perturbation-level embeddings. The module takes high-dimensional single-cell feature matrices (top), splits cells into subpopulations of interest, removes low-quality measurements and outliers, and standardizes feature distributions using non-targeting controls as reference, generating perturbation-level statistics (bottom). **d.** Batch effect correction through the Aggregate module. Violin plots show distributions of mean tubulin intensity and representative principal components across experimental batches before (top) and after (bottom) Typical Variation Normalization. **e.** Cluster module workflow. High-dimensional perturbation embeddings undergo PHATE dimensionality reduction to generate a two-dimensional representation preserving both local and global structure, followed by functional interpretation using reference database annotations, pairwise gene relationship comparisons, and group-level pathway enrichment analysis. **f.** Cluster evaluation showing precision-recall performance across Leiden resolution parameters using the CORUM benchmark. Higher resolutions yield more granular clusters with higher precision but lower recall. Precision is calculated using an adjusted metric that only counts false positives when both genes belong to different known complexes, avoiding penalization of potential novel interactions absent from reference databases. The red circle highlights resolutions 11–15, representing an optimal balance between precision and recall.”

The module supports two registration strategies: a tile-by-tile approach that matches cell patterns between corresponding tiles, and a well-level stitching approach for screens with low cell density that fail to produce a robust cell pattern at the tile level. The tile approach constructs a nine-edge hash directly from each tile, while the stitch method first stitches together the tiles into complete well mosaics using their recorded stage coordinates from microscope metadata. Once registration is performed at the full-well level, the same triangulation-based matching strategy is applied to the complete cell population, as aggregating cells across all tiles provides sufficient points for robust geometric matching in sparsely populated settings. Although the tile approach rapidly generates high-quality matches in most settings — as in this reanalysis — we applied the stitch approach to a challenging elongated human retinal pigment epithelial-1 cell dataset imaged at higher magnification. The triangle hashing approach collapsed in this setting with a median density of 19 cells per phenotyping tile, recovering only 30 match pairs. When using the stitch approach, we merged 65,480 cells out of 138,007 phenotype cells.

In our reanalysis, spatial registration between Sequencing-by-synthesis (74,406,684 cells) and Phenotype (71,629,095 cells) successfully matched and deduplicated 54,683,953 cell pairs, achieving a 76.4% recovery rate for phenotype cells and 70.4% for SBS cells. Of these matched cells, 58% (31,654,068) were successfully assigned to a single gene target, representing the effective mapping rate after spatial registration (Extended Data Fig. 5a). The approximately 25% cell loss during merge primarily reflects intentional filtering of cells at tile boundaries, where overlapping fields of view result in duplicate cell detections. This deduplication step is essential for preventing the same physical cell from contributing multiple observations to downstream analyses, and the retention rates are consistent with the expected overlap between adjacent tiles in the imaging configuration. For a representative area, the triangle hash approach achieves a 97% match rate in non-overlapping tile regions compared to 53% in overlapping regions, directly confirming that cell loss is concentrated at tile boundaries where duplicate detections are expected rather than reflecting registration failures (Extended Data Fig. 5b-c). Comprehensive visualizations enable rapid assessment of spatial distribution and quality metrics across the plate (Fig. 3b). Thus, the Merge module creates a unified dataset that directly links genetic perturbations to their corresponding phenotypic signatures at single-cell resolution.

### Classify: Cell State Classification

Analyzing cellular phenotypes across distinct cell cycle states or other biological conditions requires the ability to systematically partition cells into relevant subpopulations. The Classify module provides an integrated framework for training custom machine learning classifiers to categorize cells based on their morphological features, enabling cell-state-specific downstream analyses.

The classification workflow is designed for rapid, interactive classifier development, even for lowly represented cell populations. An interactive labeling interface rapidly displays cellular images across all wells (Extended Data Fig. 6a). For rare cell states—such as mitotic cells, which constitute only ∼4% of asynchronous cultures—feature-based gating prioritizes cells with relevant characteristics for labeling, enabling the creation of balanced training datasets without manually screening thousands of majority-class cells. The module then trains and evaluates multiple classifier architectures in parallel, automatically identifying the best-performing model. For the Vesuvius dataset, we manually annotated 800 cells (276 mitotic, 524 interphase) in less than three hours (Extended Data Fig. 6b). A high-performing model with a low number of features (XGBoost with top 50 features) achieved 96.3% accuracy on held-out training data, with strong performance for both mitotic and interphase cells (Extended Data Fig. 6c). Per-class confidence thresholds are then set interactively (Extended Data Fig 6d), and classified cells are automatically partitioned into separate analysis tracks for cell-state-specific downstream profiling. The Classify module thus enables biologically informed partitioning of heterogeneous cell populations, ensuring that downstream analyses capture cell-state-specific phenotypic variation rather than conflating distinct biological programs.

### Aggregate: From single-cell measurements to perturbation-level profiles

Optical pooled screening approaches obtain phenotype information from thousands of individual cells for each perturbation. Although this generates rich single-cell data that is useful for some applications, aggregating these measurements into perturbation-level profiles provides statistically meaningful biological insights regarding gene function. The Aggregate module transforms single-cell perturbation data into robust gene-level embeddings suitable for downstream analysis (Fig. 3c). This module allows the user to split their dataset into different subpopulations using any classifier as appropriate, before employing a standard cell feature processing pipeline with steps to filter data, align to reduce variation across experimental batches, and aggregate data to the perturbation level^19^. An optional perturbation scoring step assesses the strength of each cell’s perturbation phenotype, enabling filtering of cells with weak or noisy phenotypic effects before aggregation^20^. Batch effects arising from plate and well position are addressed through Typical Variation Normalization (TVN), which uses non-targeting control samples to estimate and remove technical variation while preserving biological signal^21^. Separately, the original interpretable features (such as nuclear size and stain intensities) are aggregated to the perturbation level to provide quantitative metrics that directly inform our understanding of which specific cellular processes drive the observed functional gene clusters. These features can be optionally bootstrapped over non-targeting control distributions to assess statistical significance, identifying which specific morphological measurements are most strongly perturbed for each gene relative to baseline variation. This enables users to move beyond aggregate profile similarity and pinpoint the individual cellular properties that drive a gene’s phenotypic signature.

A critical design choice in the aggregation pipeline is the order of dimensionality reduction and aggregation. Brieflow implements single-cell PCA, computing principal components across all individual cells before aggregating per gene. This approach captures dominant modes of cell-level phenotypic variation, so genes that push cells into similar phenotypic states have similar aggregated profiles even if their molecular mechanisms differ — emphasizing phenotypic convergence that naturally groups functionally related genes. This contrasts with an alternative approach where features are first averaged per gene before applying PCA, which instead emphasizes morphological specificity and excels at resolving fine-grained sub-pathway distinctions. Brieflow’s phenotypic convergence architecture is optimized for systematic functional discovery — identifying which genes participate in similar biological processes and placing poorly characterized genes into pathway contexts.

For the final aggregation step, cellular measurements are collapsed to the perturbation level using median aggregation, which provides robustness to outliers common in single-cell data; alternative methods including mean and trimmed mean are also supported. In the reanalysis, TVN batch correction was validated by ANOVA F-test for batch-associated variation, with p-values increasing from highly significant levels (10^-3^ for interphase) to non-significant levels (0.52 for interphase), confirming successful removal of technical variation (Fig. 3d). This produced perturbation-level embeddings for 5,299 perturbations in both interphase (30,271,668 cells; median 5,732 cells per perturbation) and mitotic (885,424 cells; median 163 cells per perturbation) populations. In sum, the Aggregate module transforms single-cell data into two complementary perturbation-level datasets: standardized principal component embeddings that serve as the foundation for clustering analysis, and aggregated interpretable feature values that enable quantitative assessment of specific cellular phenotypes for downstream biological interpretation.

### Cluster: Identification of functional relationships

The Cluster module reveals functional relationships between genes by grouping genetic perturbations with similar phenotypic profiles. This final analytical stage transforms the high-dimensional embeddings from the Aggregate module into interpretable clusters that illuminate biological pathways and functional connections (Fig. 3e).

The clustering pipeline employs PHATE^22^ (Potential of Heat-diffusion for Affinity-based Transition Embedding) for dimensionality reduction, preserving both local and global structure in a two-dimensional representation. The resulting graph weights serve as input to the Leiden community detection algorithm^23^, which identifies gene communities at a configurable resolution parameter. To objectively evaluate clustering quality, we implemented a benchmarking framework that tests gene clusters against established biological databases including STRING protein-protein interactions^24^, CORUM protein complexes^25^, and KEGG pathways^26^. Clustering is performed across a range of resolutions, and automated evaluation functions enable users to select an optimal resolution by balancing cluster granularity against enrichment for known biological relationships (Fig. 3f).

Beyond discrete cluster assignments, we quantified phenotypic divergence for each perturbation using the PHATE diffusion potential space, computing the average Euclidean distance from each gene’s potential vector to all non-targeting control vectors. These “potential distance” scores provide a continuous measure of phenotypic effect strength that complements cluster membership. Thus, the Cluster module translates high-dimensional phenotypic profiles into biologically meaningful gene groups that illuminate pathway relationships and functional connections.

For our final analysis, we selected k=12 for interphase clustering (producing 227 clusters) and k=5 for mitotic clustering (producing 222 clusters). At these resolutions, interphase clusters showed 33.0% CORUM enrichment (75 clusters), 18.1% KEGG enrichment (41 clusters), and a STRING F1 score of 0.098 (precision=0.064, recall=0.203); mitotic clusters showed 5.4% CORUM enrichment (12 clusters), 3.2% KEGG enrichment (7 clusters), and a STRING F1 score of 0.065 (precision=0.043, recall=0.128). Compared to the Funk et al. clustering at their optimal resolutions (k=10 interphase, k=9 mitotic; both producing 222 clusters), Brieflow’s interphase clustering showed a higher STRING F1 (0.098 vs. 0.067), CORUM enrichment (33.0% vs. 23.9%), and KEGG enrichment (18.1% vs. 13.5%). Mitotic clustering showed higher STRING F1 (0.065 vs. 0.044) but lower CORUM (5.4% vs. 7.2%) and KEGG enrichment (3.2% vs. 5.4%) (Extended Data Fig. 7a). The improved biological resolution relative to the original Funk et al. analysis likely reflects cumulative contributions across multiple pipeline stages: enhanced segmentation and feature extraction capture subtler morphological distinctions, and single-cell PCA preserves phenotypic variation that gene-level averaging would obscure.

### Interactive visualization and exploration dashboard

Finally, Brieflow incorporates an interactive visualization dashboard for quality control assessment and exploratory analysis (Fig. 4a). The dashboard comprises five displays — Cluster Analysis, Pipeline Stats, Quality Control, Screen Overview, and Analysis Overview — enabling users to explore gene clusters and LLM annotations, review pipeline statistics, assess data integrity, inspect experimental metadata and perturbation libraries, and examine configuration and dependencies. The dashboard additionally integrates with MozzareLLM, a large language model framework for biological interpretation described below. All finalized outputs of the reanalysis can be accessed at https://screens.wi.mit.edu/aconcagua/.

**Fig. 4.**
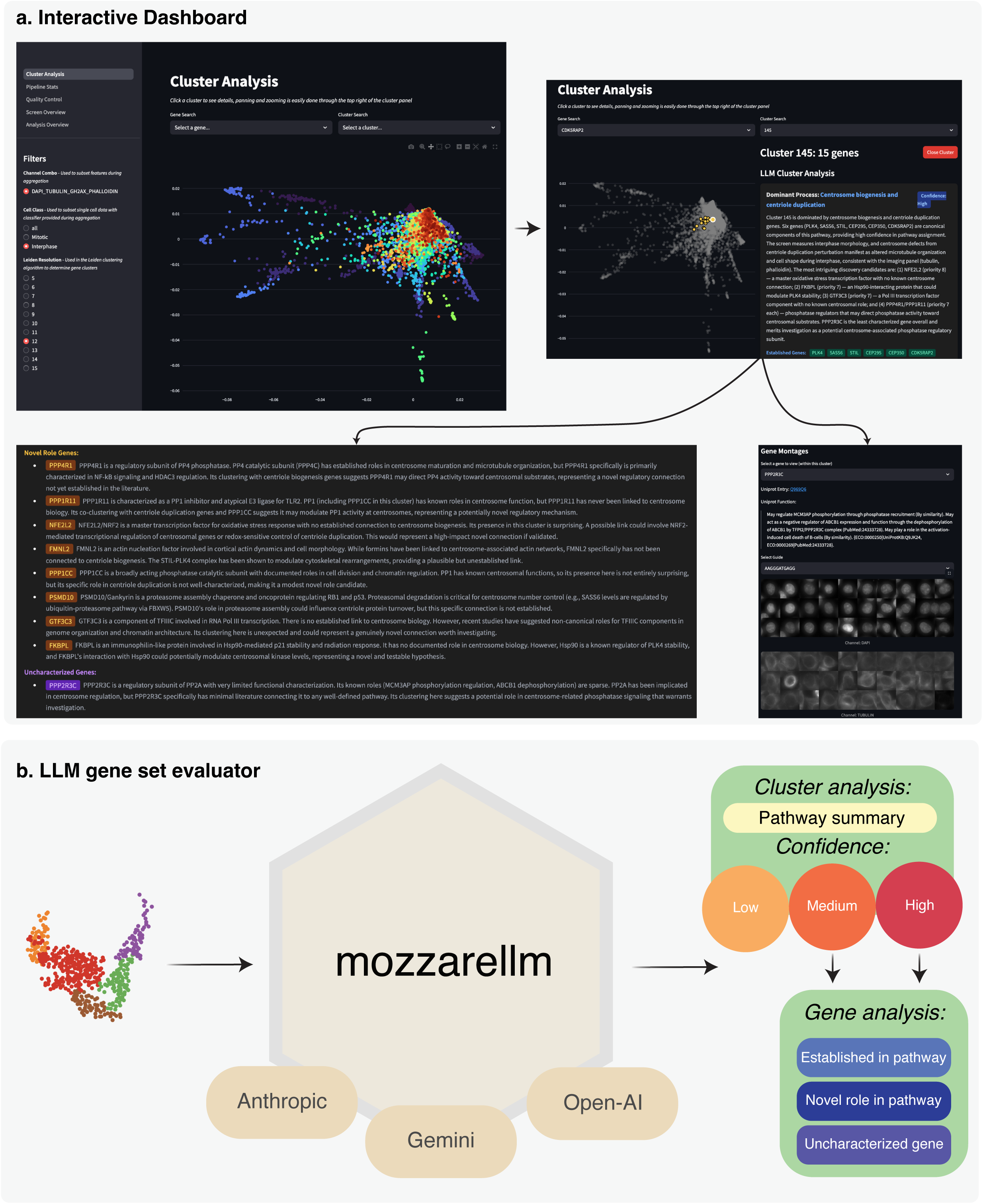
Interactive visualization dashboard and MozzareLLM for biological interpretation. **a.** Sample of the interactive visualization dashboard. The dashboard comprises five displays: Cluster Analysis for visualization of gene relationships and MozzareLLM output (displayed), Pipeline Stats for processing statistics, Quality Control for data integrity assessment, Screen Overview for experimental metadata, and Analysis Overview for configuration and dependency information. **b.** MozzareLLM workflow for automated biological interpretation of phenotypic clusters. Cluster data (left) is processed through a structured prompt engineering approach using large language models (center), producing pathway annotations with confidence levels and functional classification of constituent genes (right). MozzareLLM identifies the dominant biological process for each cluster and categorizes genes as established pathway members, characterized genes with potential novel roles, or uncharacterized genes, assigning prioritization scores based on evidence strength and discovery potential.

### MozzareLLM: Automated biological interpretation and prioritization

Although benchmarking against established biological datasets, such as CORUM and KEGG, provides confidence in cluster quality, extracting novel biological insights from these clusters remains a significant challenge relying on time-consuming and imperfect manual evaluation. Large language models (LLMs) have recently emerged as powerful tools for analyzing gene sets and interpreting biological pathways^27^, but their application to identifying novel or uncharacterized genes with potential functional significance has been limited. To address this gap, we developed MozzareLLM, a specialized LLM framework for automating the interpretation of gene-phenotype relationships and prioritizing candidates for functional investigation (Fig. 4b).

When applied to the reanalysis of the Vesuvius screen, we ran MozzareLLM separately on the clusters generated from both interphase and mitotic cell populations to capture cell-cycle-specific functional organization. For interphase clusters (n=227 at k=12), MozzareLLM identified distinct high-confidence biological processes for 73 clusters (32.2%; Supplemental Table S1), compared with 52 high-confidence clusters (23.4% of 222 clusters) from the original Funk et al. analysis at its optimal resolution (Extended Data Fig. 7b). These 73 high-confidence clusters captured 1,874 genes compared to Funk et al.’s 1,114 genes, a 68% increase in genes assigned to interpretable biological pathways. For mitotic clusters (n=222 at k=5), MozzareLLM identified high-confidence processes for 10 clusters (4.5%), capturing 731 genes compared to Funk’s 16 high-confidence mitotic clusters containing 521 genes.

To establish that MozzareLLM pathway confidence scores reflect genuine biological signal rather than artifacts of the annotation framework, we generated a shuffled negative control by randomly permuting gene-to-cluster assignments within the Brieflow interphase dataset while preserving cluster size distributions. Running MozzareLLM on this permuted clustering yielded 256 clusters, of which only 1 (0.4%) received a high-confidence pathway annotation; that single cluster contained 6 genes, and subsequent inspection revealed it arose by chance from a random co-assignment of annotated paralogs. The approximately 80-fold enrichment of high-confidence annotations in real versus shuffled data (32.2% vs. 0.4%) demonstrates that MozzareLLM’s confidence assessment is not trivially satisfied by arbitrary gene groupings; coherent pathway annotations arise only when cluster members share genuine functional relationships as reflected in their phenotypic profiles (Extended Data Fig. 7b). Consistent with this, the shuffled clustering showed 0% CORUM and KEGG enrichment (compared to 33.0% and 18.1% for real interphase clusters) and a STRING F1 score of 0.003 (compared to 0.098), confirming that the biological signal captured by Brieflow’s clustering is absent in randomized controls. High-confidence Brieflow clusters contained, on average, 16.6 genes with established functions in the identified pathway, 7.4 genes with established functions but potential novel roles in the identified processes, and 1.7 genes with limited functional annotation — representing candidates that co-cluster with well-characterized pathway components (Extended Data Fig 7c, Supplemental Table S2).

### Reanalysis with Brieflow recovers mitochondrial clusters

A powerful feature of optical pooled screening is its ability to report on diverse cellular properties regardless of the markers used, as core cellular processes are intimately interrelated. Despite the use of only four cellular markers — DNA, a DNA damage marker, actin, and microtubules — both the original Funk et al. analysis and our reanalysis detected coherent gene clusters for protein translation, protein degradation, vesicle trafficking, and numerous other cellular processes.

A notable absence from both the initial Funk et al. analysis relative to complementary Perturb-seq analyses^28^ were gene clusters corresponding to mitochondrial function. Mitochondrial biology provides a demanding test case due to the absence of imaging channels in the Vesuvius screen that directly stained mitochondria, such that any signal must arise from indirect morphological consequences of mitochondrial perturbation. MozzareLLM reanalysis of the Funk et al. clustering identified four interphase clusters containing groups of mitochondrial proteins (Fig. 5b), with established-gene fractions ranging from 47.4% to 71.4% and mixed functional content throughout. However, these clusters were missing the vast majority of established mitochondrial components and contained non-mitochondrial genes forming coherent sub-programs rather than isolated outliers. For example, Funk interphase cluster 149 (19 genes, 52.6% established) co-clusters the RAF-MAPK kinases KRAS and BRAF with its OXPHOS core, and Funk interphase cluster 147 (19 genes, 47.4% established) embeds an intact actin disassembly module (CAP1, CFL1, WDR1) alongside genuine OXPHOS subunits.

**Fig. 5.**
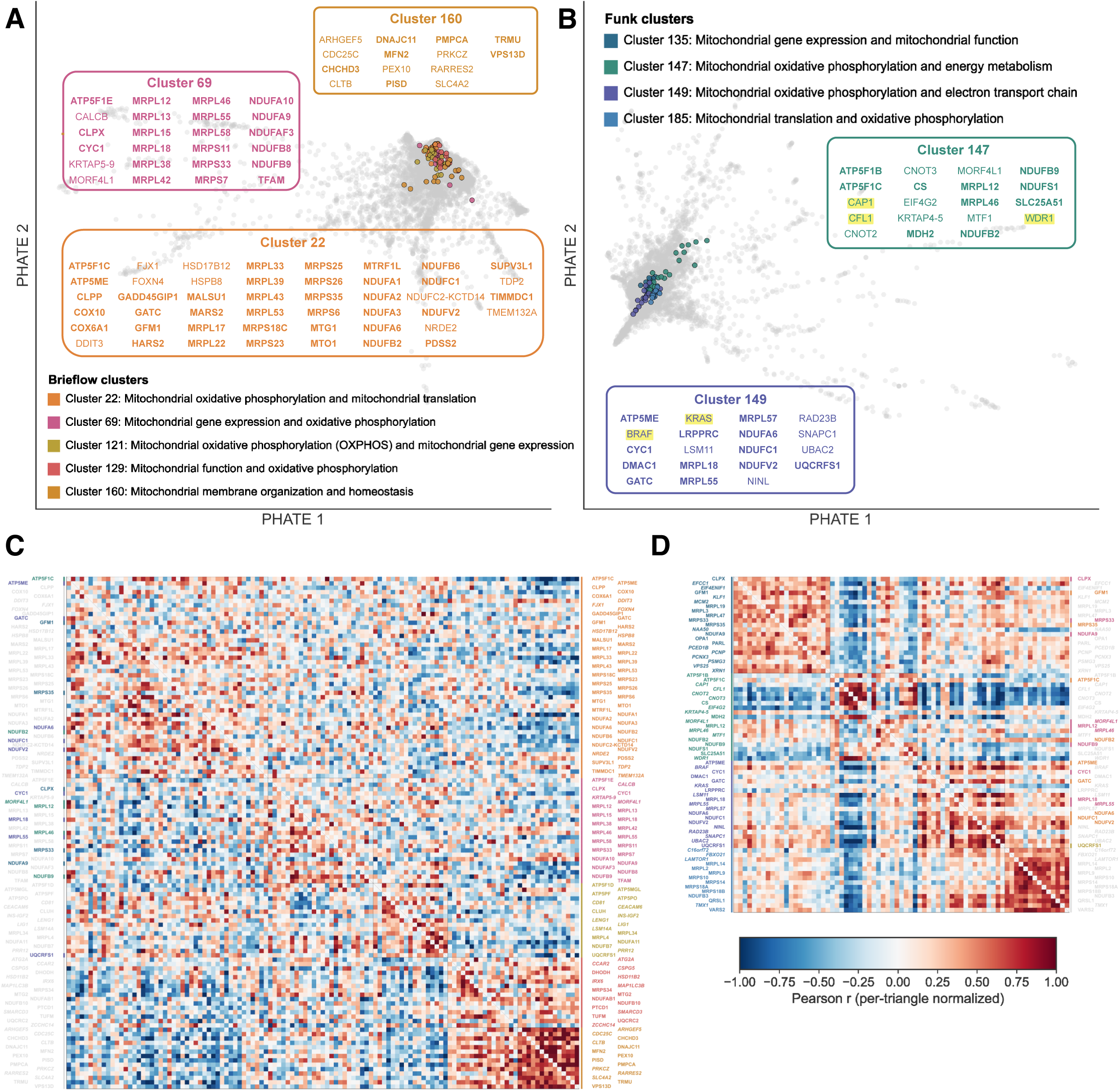
Recovery of mitochondrial sub-programs by Brieflow. **a.** PHATE visualization of Brieflow interphase clustering with five high-confidence mitochondrial sub-modules highlighted. Labeled mitochondrial submodules with bold denoting validation via MitoCarta3.0 **b.** Similar PHATE visualization from the Funk et al. analysis with four mitochondrial-containing clusters highlighted. **c.** Pearson correlation matrix of PCA-based perturbation profiles (Brieflow’s primary aggregation methodology) for mitochondrial genes, organized by Brieflow cluster assignments. Gene labels are colored by cluster membership as in **a**; bold labels indicate established pathway genes, italic labels indicate uncharacterized or novel-role genes. **d.** Pearson correlation matrix of raw feature-based perturbation profiles (Funk et al.’s primary aggregation methodology) for the same genes, organized by Funk et al. cluster assignments. Gene labels are colored by cluster membership as in **b**, with the same bold/italic convention.

In contrast, MozzareLLM applied to Brieflow clusters highlights five distinct high-confidence mitochondrial sub-modules (Fig. 5a), encompassing 119 genes of which 81 (68.1%) are MitoCarta-annotated^29^—nearly twice the number of mitochondrial genes recovered by the Funk et al. analysis at higher purity. Brieflow interphase cluster 22 (48 genes, 79.2% established) covers OXPHOS assembly and mitochondrial translation; cluster 69 (24 genes, 87.5% established) concentrates mitochondrial ribosome proteins and gene expression factors including TFAM; clusters 121 and 129 resolve ATP synthase assembly and mitochondrial translation machinery, respectively; and cluster 160 captures mitochondrial membrane organization (Fig. 5c). The few novel-role genes present appear as isolated individuals with plausible mitochondrial connections—such as the ISR effector DDIT3 and the mitophagy chaperone HSPB8—rather than as coherent non-mitochondrial sub-programs. Together, clusters 22 and 69 alone account for 24 distinct MRPL/MRPS mitochondrial ribosome subunits distributed across just two clusters, compared with the scattering of mitoribosomal proteins across multiple diverse clusters in the Funk et al. analysis. Most distinctively, cluster 160 captures a membrane organization sub-program — MFN2 (fusion), DNAJC11 (cristae architecture), PISD (phospholipid synthesis), VPS13D (ER–mitochondria contacts) — with none of its established genes appearing in any Funk mitochondrial cluster, representing sub-compartment specificity that the original analysis did not recover (Fig. 5c). Correlation of raw interpretable features between the two datasets (Fig. 5d), corresponding to the Funk et al. aggregation approach, shows relatively similar distributions for these clusters—likely because the large number of cells per perturbation produces stable feature means. In contrast, correlation of PCA-based embeddings (Fig. 5c), corresponding to the Brieflow aggregation approach, reveals more defined differences between the strategies, suggesting that computing principal components at the single-cell level before aggregation preserves phenotypic variation that gene-level feature averaging may obscure.

To assess whether Brieflow retains the principal biological findings of Funk et al., we tracked all 57 clusters highlighted in that study as biologically interpretable, 15 of which were discussed individually in the text as exemplary functional modules and the remaining 42 organized into pathway groups and presented as heatmaps or annotated on PHATE plots. Using MozzareLLM, we independently assigned high-confidence pathway annotations to 50 of 57 (87.7%) of the original Funk clusters. In the Brieflow clustering, 77.2% of these curated clusters were at least partially preserved (≥50% of genes co-occurring in a single Brieflow cluster), with 26 (45.6%) well-preserved at a mean retention of 91.8%. Among the 15 highlighted clusters, 7 were well-preserved and the remaining 8 showed partial preservation or fragmentation (Extended Data Fig. 8a); however, 82% of the redistributed genes from these 8 partially preserved clusters were assigned to high-confidence Brieflow modules (Extended Data Fig. 8b), indicating that gene redistribution typically reflects reassignment into more specific modules rather than loss of signal.

Beyond recapitulating the findings of Funk et al., we next asked whether Brieflow resolves functional programs not captured by the original analysis. We computed bidirectional Jaccard similarity indices between all high-confidence clusters across both interphase and mitotic populations, defining low concordance as Jaccard < 0.15 (Supplemental Table S4). This revealed a pronounced asymmetry: 29 of 83 high-confidence Brieflow clusters (35%) had no adequate match in the Funk clustering, compared with 14 of 68 Funk clusters (21%) that were reciprocally unmatched (Extended Data Fig. 9a, b). Notably, 297 genes in Brieflow-unique clusters were flagged as novel-role or uncharacterized, representing candidates whose functional assignments are potential targets for follow-up investigation.

## Discussion

Although technologies such as Perturb-seq^30^ have benefited from extensive methodological refinement through the widespread use of gene expression data, optical pooled screening represents an emerging data type with substantial room for methodological innovation compared with more established functional genomics approaches. The unique analytical challenges inherent to OPS—including multi-modal data integration across different imaging modalities, complex image processing requirements, and the extraction of biologically meaningful features from high-dimensional cellular measurements—necessitate specialized computational approaches that continue to evolve rapidly. These challenges present significant opportunities for innovation in computational methods tailored specifically to the complexities of image-based functional genomics.

Brieflow represents a major advancement in the analysis of OPS data, addressing key bottlenecks that have limited the widespread adoption of this powerful technique. Although experimental OPS platforms have become increasingly accessible, challenges in analysis represent a significant barrier to entry for biologists seeking to adopt OPS, underscoring the need for accessible, integrated computational tools that can bridge the gap between experimental accessibility and analytical complexity. Although existing OPS analysis approaches have made valuable contributions, they often focus on specific analytical challenges rather than providing integrated and validated end-to-end workflows, with some relying on specialized software environments that limit scalability and broader adoption (Table 1). By providing a unified framework for end-to-end analysis, Brieflow eliminates the need for fragmented pipelines and format conversions, substantially reducing the technical barriers to entry for researchers interested in applying OPS to their biological questions, while its modular architecture allows users to adapt individual components to their specific experimental contexts. Our reanalysis of the large-scale ‘Vesuvius’ optical pooled screen demonstrates Brieflow’s capabilities in handling large-scale datasets to reveal enhanced biological insights while maintaining high data quality and analytical rigor.

**Table 1:** Comparison of existing optical pooled screening analysis pipelines and their capabilities. Features are compared across four computational frameworks for OPS data analysis. Periscope relies on CellProfiler-based architecture requiring integration of multiple separate analytical pipelines. OpticalPooledScreens provides basic phenotype and sequencing analysis components but lacks comprehensive integration and documentation for end-to-end workflows. PerturbView focuses specifically on tissue spatial alignment applications with limited broader analytical scope. Brieflow provides the first unified framework encompassing complete analysis from raw data to biological interpretation. Extensive benchmarking of alternative pipelines was not feasible due to limited accessibility and the time-intensive nature of adapting existing frameworks to perform the comprehensive end-to-end analysis demonstrated with Brieflow.

| Feature | PERISCOPE<sup>3</sup> | OpticalPooledScreens<sup>9</sup> | PerturbView<sup>13</sup> | Brieflow |
| --- | --- | --- | --- | --- |
| End-to-End Integration | ? | ? | ? | ? |
| Unified Framework | ? | ? | ? | ? |
| Analysis Documentation | Comprehensive | Minimal | Minimal | Comprehensive |
| Workflow Management | Manual | Snakemake | Snakemake | Snakemake |
| Modular Architecture | ? | ? | ? | ? |
| Community Extensibility | ? | ? | ? | ? |
| Primary Focus | CellProfiler-based analysis | Basic phenotype/SBS | Tissue spatial alignment | Complete OPS analysis |
| Data Integration | Multiple separate pipelines | Fragmented workflow | Specialized use case | Single integrated pipeline |
| Analytical Scope | Complete analysis, no interpretation | Incomplete pipeline | Specialized application | Complete analysis and interpretation |

The standardization and computational reproducibility provided by Brieflow will be particularly important as the adoption of OPS approaches continues. Through Snakemake-based workflow management and comprehensive configuration documentation, the Brieflow pipeline enables researchers to share not only their results but also the exact methodologies used to generate them, enhancing transparency and facilitating collaborative research. The open-source implementation of Brieflow, with its well-documented interfaces and extensible design, provides a foundation for community-driven improvement and customization. The modular architecture enables rapid development of new capabilities—for example, swapping in a custom segmentation method or extending feature extraction to novel cellular structures can be accomplished rapidly while maintaining the pipeline’s organization and validation framework. As with any complex analytical pipeline, certain implementation choices reflect practical constraints that users should consider.

Although Brieflow provides a comprehensive framework for optical pooled screening analysis, several technical considerations should be noted when applying the pipeline to new experimental contexts. The current implementation assumes one-to-one tile correspondence within phenotyping rounds and sequencing-by-synthesis cycles, a constraint satisfied by most standard microscope configurations. Notably, despite images being acquired on two different microscopes for the SBS and phenotyping workflows in our validation dataset, the merge registration performs robustly, demonstrating that the pipeline can successfully integrate data across imaging platforms when tile layouts are consistent.

Registration quality depends on the ability to identify corresponding cellular landmarks across imaging rounds. The default tile-by-tile registration performs robustly under standard screening conditions, but high-magnification screens with sparse cellular populations may not provide sufficient landmarks per tile for robust geometric matching. In these cases, Brieflow’s well-level stitching strategy aggregates cells across tiles to recover sufficient match points, as demonstrated in our retinal pigment epithelial cell dataset. Conversely, the stitching approach introduces greater computational demands during pairwise distance matrix construction for screens with large numbers of cells per well. These trade-offs are being actively addressed through ongoing improvements to Brieflow’s registration algorithms and memory handling.

Our CellProfiler-inspired feature extraction, although effective for morphological profiling, reflects a broader challenge in the field: reliable methods to extract single-cell image features in a fully Pythonic format are not widely available. As the field of single-cell image featurization continues to advance with improved deep learning-based approaches and standardized feature representations, Brieflow’s modular architecture is designed to readily incorporate these improvements.

Brieflow is in active development and will remain so for the foreseeable future. The pipeline benefits from contributions from an expanding user community, and improvements to individual modules—from segmentation to feature extraction to pathway analysis—will continue to enhance the quality of biological insights derived from optical pooled screens. A central priority is aligning Brieflow with next-generation file formats (NGFF), particularly OME-Zarr, to improve interoperability with the broader imaging ecosystem and enable more efficient processing of increasingly large datasets.

The pipeline’s modular architecture positions it as a central hub for OPS analysis, where advances in component methods can be seamlessly integrated to benefit the entire workflow. Furthermore, the standardized outputs produced through Brieflow constitute valuable resources for the development of machine learning models on functional genomics data. As perturbational datasets capture a broader spectrum of cellular phenotypes, the analysis of multiple OPS experiments processed through a common framework enables the creation of comprehensive training datasets with enhanced generalizability and predictive power. These standardized datasets will significantly advance computational cell modeling efforts, allowing researchers to build increasingly accurate in silico predictions of cellular behaviors that integrate multi-dimensional phenotypic data across diverse genetic perturbations.

Finally, the addition of MozzareLLM for automated biological interpretation represents a complementary approach to extracting actionable insights from high-dimensional phenotypic data. By leveraging large language models specifically oriented towards biological knowledge extraction, this component accelerates the transition from data to hypothesis formation, a critical step in scientific discovery of gene function.

## Code Availability

All code is publicly available. The core Brieflow pipeline is available via GitHub at https://github.com/cheeseman-lab/brieflow. The specific implementation for the HeLa OPS dataset analysis described in this paper, including benchmarking experiments between methods, is available at https://github.com/cheeseman-lab/aconcagua-analysis. MozzareLLM, designed to scale to different types of gene grouping analyses beyond its integration with Brieflow, is available as a standalone tool at https://github.com/cheeseman-lab/mozzarellm. Comprehensive documentation for Brieflow, including tutorial videos and detailed implementation instructions, can be accessed at https://brieflow.readthedocs.io. For researchers interested in analyzing optical pooled screening data, we recommend consulting this documentation before initiating a new analysis project based on the Brieflow-analysis template. Any questions, issues, or code bugs should be reported at https://github.com/cheeseman-lab/brieflow/issues. A small test image dataset for validating Brieflow installation and testing pipeline functionality is included in the Brieflow repository at https://github.com/cheeseman-lab/brieflow/tests/small_test_analysis. This dataset provides representative examples of OPS images for both sequencing-by-synthesis and phenotyping outputs. All benchmarking scripts conducted as part of this study are also available at https://github.com/cheeseman-lab/aconcagua-analysis/benchmarks.

## Data Availability

The raw dataset (>20TB) used in this study can be provided upon reasonable request by contacting the corresponding authors, as sharing raw imaging data of that size is a considerable challenge. A stable interactive interface for exploring the results of the HeLa screen analysis, including visualizations of phenotypic clusters and gene relationships, quality control plots, details on the Brieflow run conducted, and experimental metadata, is publicly accessible at http://screens.wi.mit.edu/aconcagua, from which cluster groupings and feature matrices can be downloaded.

## Acknowledgements

This work was supported by grants from the NIH (R35GM126930 to I.M.C., R01HG009283 to P.C.B.), and the Chan Zuckerberg Initiative to I.M.C. and P.C.B. M.D. is supported in part by an NSF GRFP fellowship. This research was also supported by the Whitehead Innovation Initiative and the Massachusetts Life Science Center Data Science Internship Program. A.M. was supported by the MIT Undergraduate Research Opportunities Program. We extend our gratitude to David Feldman for the original implementation of the pipeline. We thank Luke Funk for useful information on how the screen was conducted and in-depth discussions on how it was analyzed, and both Luke Funk and Kuan Chung Su for conducting the HeLa screen. We acknowledge members of the Blainey and Cheeseman laboratories for their valuable discussions regarding pipeline development, and especially Russell Walton for his guidance and for providing the secondary ISS dataset for additional benchmarking. We are particularly grateful to members of the Whitehead Institute Information Technology Core for their assistance with pipeline acceleration and implementation on high-performance computing infrastructure. <u>We thank Charlie Bushman of Sunbeam for his guidance on using Snakemake to build adaptable pipelines for biological data.</u> We also thank Alán F. Munoz and members of the Broad Imaging Platform for their insights regarding CellProfiler implementation. Additionally, we acknowledge Albert Dominguez Mantes for his thoughtful discussions concerning the implementation of Spotiflow.

## Author contributions

M.D. and R.K. conceptualized the work. M.D., R.K., A.K.C.D., A.M., S.J.C. and A.N-U. implemented the software. M.D., I.C., and P.C.B. drafted the manuscript. All co-authors reviewed and edited the final manuscript.

## Methods

### System Design and Implementation Architecture

Brieflow follows a standardized Snakemake project development template with four principal components: library code containing core analytical functions, Python scripts for individual rules, Snakemake rules defining execution logic, and targets specifying pipeline outputs. This architecture separates core analytical logic from execution scripts, enabling modification of specific components without disrupting the overall pipeline.

The data structure employs standardized file formats including TIFF for image data, TSV and parquet for tabular data, and PNG for evaluation outputs. Consistent naming conventions facilitate automated data organization and rule matching, while clearly separating raw, intermediate, and analysis data.

Brieflow-analysis, provided as a separate template repository, incorporates a hierarchical YAML-based configuration system organized in a tree structure that separates experimental parameters from processing logic. Users create a new repository from this template for each screen analysis, with Brieflow integrated as a git submodule to ensure consistent versioning while enabling project-specific customization. The repository includes comprehensive testing procedures to verify correct installation across computing environments. Brieflow implements multi-level parallelization across sample, plate, well, and tile levels, with GPU acceleration available for computationally intensive tasks such as deep learning-based segmentation. The framework includes specialized execution scripts that optimize job submission for different computing environments, processing plates in sequence to avoid overwhelming computational resources and consolidating multiple small tasks into fewer, larger jobs. The system includes configuration profiles for different execution environments that specify resource allocation, job submission parameters, and environment variables, and makes use of the Snakemake plugin catalog for use of Brieflow across different user-specific interfaces.

The framework supports interchangeable components through specialized branches designed for different experimental contexts, enabling community contributions in specific areas such as spot calling, segmentation, or feature extraction. Standardized interfaces between modules ensure that modifications to specific components can be implemented without disrupting the overall workflow.

### Preprocess Module

The Preprocess module converts raw microscopy files to standardized TIFF format while extracting comprehensive metadata including spatial information, optical parameters, and experimental identifiers. The module employs a uniform file reading architecture that supports multiple input formats through a configurable interface: users specify their file type (ND2 or TIFF) and associated metadata structure, and the pipeline automatically adapts its reading and parsing logic accordingly. This design enables compatibility with diverse microscopy systems including Nikon (ND2), Phenix (TIFF), and Squid (TIFF) platforms.

The module processes diverse file organizations including well-based image storage or individual tile-based acquisitions, combined or separated imaging rounds, and single-channel or multi-channel file structures. Brieflow also supports multiple-round phenotyping via iterative staining, with each round treated as an independent phenotyping cycle that is processed and aligned separately before downstream integration. The current implementation assumes one-to-one tile correspondence within phenotyping rounds and sequencing-by-synthesis cycles, which is satisfied by most standard microscope configurations.

Configuration parameters are set through interactive notebooks where users specify their input files and file format type. For ND2 files, metadata is extracted directly from the proprietary format; for TIFF files, users provide accompanying metadata files or configure parsing rules for filename-encoded information (e.g., plate, well, tile, and channel identifiers). This automatically generates a comprehensive file mapping all combinations of plates, wells, tiles, and channels, eliminating the need for hardcoded values in downstream analysis. Illumination correction is implemented through batch-level correction field calculation, computing illumination patterns by averaging multiple fields of view across an experimental batch. Users can specify a sample fraction (ranging from 0-1) to control the proportion of images used for correction field estimation, with our implementation using the complete dataset. After smoothing with a median filter, each image undergoes correction by division with the illumination function in downstream modules. The implementation leverages multi-threading capabilities to accelerate processing of large image datasets.

### Sequencing By Synthesis Module

The Sequencing-by-synthesis module processes *in situ* sequencing images through a sequential workflow: alignment of sequencing cycles, illumination correction optimized for sequencing signal detection, application of a Laplacian-of-Gaussian filter to enhance spots, and computation of intensity standard deviation to identify consistent signals across cycles.

Alignment supports multiple methods, including reference-based alignment using DAPI as a fiduciary marker (if imaged in each round) and signal-based alignment using the mean sequence signal. Alignment is performed using phase cross-correlation with a configurable upsampling factor for sub-pixel precision; our implementation used integer-pixel resolution, which was sufficient. To account for sub-pixel alignment errors, the module applies maximum filtering with a configurable width (set to 3 pixels in our analysis) that dilates sequencing signals slightly, ensuring robust signal capture despite minor registration imperfections.

For barcode detection, two distinct approaches are implemented: a standard method based on statistical signal detection and a deep learning-based method using Spotiflow. The standard approach (which we implemented with a peak threshold of 400) first applies a Laplacian-of-Gaussian (LoG) filter (with kernel width σ = 1 pixel) to enhance spots across sequencing channels. After applying this filter to all sequencing cycles, the standard deviation across cycles and mean across channels are computed to identify consistent signals. Local maxima are then identified within a defined neighborhood (default width of 5 pixels) using maximum and minimum filters: at each local maximum, the peak score is calculated as the difference between the maximum and minimum values within the neighborhood. Peaks within the neighborhood width of image borders are excluded to avoid edge artifacts. This approach effectively captures spots that maintain consistent signal across multiple sequencing cycles while filtering out noise and non-specific background. The Spotiflow method provides an alternative approach, utilizing a pre-trained U-Net model to predict spot locations from a selected sequencing cycle across all four base channels. Each base channel is processed independently, and results are combined while enforcing minimum distance constraints through iterative distance-based filtering to exclude overlapping signals. For benchmarks, Spotiflow was run with a probability threshold of 0.3, using the first cycle for spot detection, and a minimum distance of 1 pixel between detected spots.

Base calling implements a statistical approach that considers channel cross-talk and signal-to-noise ratios. Each called base receives a quality score reflecting confidence in the call, calculated as:

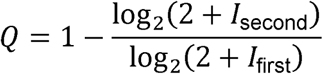

where *I*_first_ and *I*_second_ are the intensities of the highest and second-highest channels, respectively. The module corrects for varying base intensities using either median-based or percentile-based normalization (we selected the median-based approach, based on better correction of A-C crosstalk). For median-based correction, a transformation matrix is constructed by: (1) for each channel, identifying spots where that channel has maximum intensity; (2) computing the median intensity profile across all channels for these spots; (3) assembling these profiles into a matrix and normalizing columns to sum to 1; and (4) inverting this matrix to obtain the correction transformation. For percentile-based correction, the 95th percentile of relative intensities (intensity divided by row sum) is used instead of the median to identify high-confidence spots for each channel. Both methods support restriction of correction estimation to reads within cells, thereby excluding aberrant signals.

Following base calling, reads are mapped to a reference barcode library to identify genetic perturbations. The mapping process supports both exact matching and error-tolerant matching with configurable edit distance thresholds. For error correction, two complementary approaches are implemented: (1) reference-based correction, which maps reads to the nearest barcode in the reference library within a specified Hamming distance threshold, resolving ambiguous mappings by selecting the barcode with the smallest edit distance or flagging reads with multiple equidistant matches; and (2) quality-weighted correction, which incorporates per-base quality scores to prioritize high-confidence base calls when resolving mismatches. The module also supports combinatorial barcode schemes used in PerturbView and Zombie *in situ* sequencing approaches, where multiple barcode segments are read and concatenated to form a complete perturbation identifier. For these multi-segment barcodes, each segment is called independently, and segments are then recombined according to user-specified rules to reconstruct the full barcode sequence before mapping to the reference library. Segmentation of cells in SBS images is performed after illumination correction, with support for Cellpose, MicroSAM, StarDist, and Watershed; we employed Cellpose with the cyto3 model (nuclei diameter 9.44 pixels, cell diameter 18.85 pixels) using base C of the last sequencing cycle due to high background signal in other channels. Segmentation masks are passed to base extraction, which integrates intensities at detected peak locations restricted to either the cellular or nuclear compartment as configured — the nuclear compartment option directly supports T7-based and Zombie *in situ* sequencing methods where amplified DNA barcodes are confined to the nucleus.

Barcode-to-cell assignment resolves a single perturbation identity per cell from the full set of mapped reads. A critical parameter governing this step is the barcode prioritization strategy, which can be set to either peak- or count-prioritized calling. Peak-prioritized calling selects the barcode associated with the highest-intensity spot and is the appropriate mode for T7-based and Zombie *in situ* sequencing chemistries, where each cell produces a single bright amplified locus. Count-prioritized calling instead selects the most frequently observed barcode identity across all reads within a cell and is appropriate for mRNA-based barcoding approaches where multiple distinct transcript species from different loci are detected per cell. The two modes produce mutually exclusive output metrics — peak intensity scores in peak mode, read counts in count mode — and selecting the correct mode for the sequencing chemistry used is essential for accurate perturbation assignment. For our primary analysis, we used count-prioritized calling.

The module is designed to accommodate diverse *in situ* sequencing chemistries through configurable parameters including spot detection area, peak detection width, and intensity thresholds. To facilitate parameter optimization before full-scale processing, a grid search utility systematically evaluates combinations of detection parameters on a representative subset of tiles, reporting spot count, mapping rate, and cell assignment rate for each combination.

To validate the SBS module’s generalizability, we applied it to an independent *in situ* sequencing dataset with three known barcode constructs sequenced via T7-based chemistry across three cycles. We used the standard spot detection method with a peak threshold of 400, a peak width of 5, peak-prioritized barcode calling, and reads assigned to the nuclear compartment, consistent with T7-based chemistry where amplified barcodes localize to the nucleus.

### Phenotype Module

The Phenotype module extracts cellular measurements through multiple processing steps: specimen-specific illumination correction, alignment of fluorescence channels to correct for chromatic aberrations, and segmentation of nuclear and cellular boundaries. The implementation supports both single-step and multi-step alignment strategies, with options for custom offsets when specific channels require targeted adjustment.

In alignment with previous methodologies, phenotype images were first maximum intensity projected to compress z-slices into a single plane. Channel alignment is performed using phase cross-correlation with a configurable upsampling factor (default of 2) for sub-pixel precision. A centered window of the image data can be used for alignment calculation to reduce noise from image edges. The computed offsets are applied via similarity transformations to shift channels into alignment.

At high magnifications or when imaging multiple phenotyping rounds with iterative staining protocols, automated alignment may be insufficient due to larger inter-round shifts or differences in sample positioning. For these cases, the module supports a two-stage manual alignment approach: first, users specify coarse-grain offsets (in pixels) to bring rounds into approximate registration based on visual inspection of landmark features; second, the automated phase cross-correlation refinement is applied to achieve precise sub-pixel alignment. This hierarchical strategy enables robust alignment even when initial offsets exceed the capture range of correlation-based methods. Custom channel-specific offsets can also be specified when particular channels exhibit systematic chromatic aberrations that differ from the global alignment solution.

For cell segmentation, the module implements a primary workflow to identify nuclear and cellular boundaries, followed by a derived segmentation step that extracts cytoplasmic regions by subtracting nuclear masks from cellular masks. Multiple segmentation approaches are supported, including deep learning-based methods such as Cellpose, MicroSAM, and StarDist, each optimized for different cellular morphologies and imaging conditions. Our implementation used Cellpose with the cyto3 model, with nuclei diameter of 42.68 pixels and cell diameter of 68.06 pixels, and default threshold parameters.

Brieflow supports two configurable feature extraction approaches: a CellProfiler-inspired pure Python emulator developed as part of Brieflow, and the cp_measure package, a pure Python CellProfiler implementation developed by the CellProfiler team. The Brieflow emulator was developed to address integration challenges between Java-based CellProfiler and Brieflow’s Snakemake-based workflow architecture, and integrates algorithms from scikit-image, mahotas, and SciPy. The emulator extracts four categories of features. Intensity features include integrated intensity, mean, standard deviation, maximum, minimum, edge intensities (using morphological operations with connectivity of 2), mass displacement, quartiles, median, and median absolute deviation. Texture features comprise Haralick features computed via mahotas and PFTAS (Parameter-Free Threshold Adjacency Statistics). Shape features include Zernike moments, Feret diameters, Hu moments, convex hull metrics, eccentricity, and solidity. Correlation and colocalization features include Pearson correlation between channel pairs, least-squares slope, overlap coefficient, Manders’ coefficients (M1 and M2), and rank-weighted colocalization coefficients, computed using Otsu thresholding. For user-specified channels, specialized foci detection and quantification is implemented using a white tophat filter followed by Laplacian-of-Gaussian enhancement to enable counting and characterization of subcellular punctate structures, which we configured for the GH2AX channel (index 2). Detected foci are refined using watershed segmentation on the distance transform of the thresholded image. Feature extraction operates at three distinct cellular compartments—nucleus, cell, and cytoplasm—allowing precise characterization at each level. For spatial relationships between cells, the module calculates neighbor metrics including counts of adjacent cells and percent boundary contact. A complete description of all extracted features and their definitions is provided in the Brieflow repository. Brieflow’s modular architecture is designed to accommodate future integration of improved feature extraction methods as they mature.

### Merge Module

The Merge module integrates data from Sequencing by synthesis and Phenotype modules through spatial registration between imaging modalities. The foundation of the merge process is accurate alignment that identifies corresponding fields of view between datasets, leveraging metadata from the Preprocess module.

The implementation employs a Delaunay triangulation strategy to create hash-based descriptors for cell patterns, enabling robust matching even with partial cell correspondence. For each tile, a Delaunay triangulation is computed from cell centroid coordinates, requiring at least 4 valid cells per tile. For each triangle in the triangulation (excluding those on the outer boundary), a nine-edge hash descriptor is computed: this includes the three internal edges of the triangle plus six edges connecting to the three neighboring triangles. The edges are ordered by starting from the longest internal edge to ensure consistent orientation. Each hash consists of the 18-dimensional vector of edge displacements (9 edges × 2 coordinates), along with the triangle center computed as the mean of vertex coordinates. The nine-edge hash method extracts vector displacements for edges connected to each triangle, creating a distinctive signature that can be matched across imaging modalities despite differences in magnification and optical characteristics.

The merge process uses a multi-step alignment algorithm that iteratively refines the transformation model as more matching points are identified. In the tile-by-tile approach, initial alignment begins with a small set of candidate tile-site pairs, either specified by the user or automatically determined. Our implementation used six predefined tile pairs, providing starting points for the iterative alignment. For each candidate pair, the algorithm evaluates the match by: (1) computing normalized edge vectors (dividing by magnitude); (2) finding nearest neighbors between vector sets using squared Euclidean distance; (3) filtering triangle matches below a distance threshold (default 0.3); (4) fitting a RANSAC regressor to the matched triangle centers to estimate the affine transformation; and (5) computing a match score as the proportion of transformed triangle centers within a point threshold (default 2 pixels) of their nearest neighbors.

To control alignment quality, the pipeline offers configurable parameters for determinant range, score threshold, and distance threshold. Our implementation used a determinant range of [0.06, 0.065], ensuring the scaling factor remained within acceptable bounds. The matching score threshold was set to 0.1, requiring a minimum level of triangle pattern overlap to consider a match valid. For cell-level matching, a distance threshold of 2 pixels controlled the maximum distance between transformed coordinates for cells to be considered matches.

The iterative alignment proceeds by: (1) filtering matches by determinant range and score threshold; (2) using RANSAC regression on matched tile-site coordinates to predict additional candidate pairs; (3) evaluating new candidates in parallel batches (default batch size of 180); and (4) repeating until no new matches are found. This approach progressively expands from initial known matches to identify all valid tile-site correspondences.

With spatial registration established, individual cells between modalities are linked based on their spatial coordinates, applying the optimized transformation parameters. The matching process incorporates distance-based thresholds to filter low-confidence associations. A deduplication pipeline resolves cases where multiple cells from one modality match a single cell in the other modality, applying a hierarchical strategy that first selects the best Sequencing by synthesis match for each phenotype cell (based on whether a cell has a known barcode), then selects the best phenotype match for each remaining Sequencing by synthesis cell (based on highest minimum channel intensity).

Quality assessment evaluates alignment quality using determinant range checks to verify scaling consistency and scoring parameters to assess matching accuracy. After cell matching, match rates for both modalities are calculated, reporting the percentage of cells successfully integrated into the final dataset.

For experimental conditions where cell density is low or tiles contain insufficient cells for robust nine-edge hash construction (which requires at least 4 cells per tile for Delaunay triangulation), the module implements an alternative well-level stitching approach. In this method, individual tiles from each imaging modality are first stitched together into complete well mosaics using their recorded stage coordinates from microscope metadata. The stitching process assembles tiles into complete well mosaics according to their recorded stage positions, with optional blending at tile boundaries to reduce edge artifacts. Once both modalities are reconstructed as full-well images, registration is performed at the well level rather than tile level, using the same triangulation-based matching strategy but applied to the complete cell population within each well. This approach aggregates cells across all tiles, providing sufficient points for robust geometric matching even when individual tiles are sparsely populated. The well-level alignment transformation is then applied to map cell coordinates between modalities. This stitching-based strategy is particularly valuable for screens with low cell confluence, small tile sizes, or high cell loss during sample preparation, where tile-by-tile matching would otherwise fail due to inadequate cell numbers for triangulation.

### Classify Module

The Classify module implements a complete machine learning pipeline for training and applying custom cell classifiers. The implementation is organized into four main components: labeling, training, calibration, and application.

The interactive labeling system presents cellular images in batches, with each batch containing a mixture of in-gate cells (prioritized by user-specified feature thresholds) and out-of-gate cells (randomly sampled for diversity). For our implementation, we configured the interface to display 25 cells per batch with 1 out-of-gate example, using DAPI and Tubulin channels for visualization. Feature gating prioritized cells with high nuclear DAPI median absolute deviation (90th percentile and above) to efficiently identify mitotic cells with condensed chromatin. The labeling interface supports incremental dataset expansion, loading previously labeled examples and filtering them from the candidate pool to avoid redundant annotations. Users can optionally enable relabeling mode to revisit and modify existing annotations. Labeled cells are continuously saved to a checkpoint file to prevent data loss during annotation sessions, with final training datasets exported as parquet files containing both morphological features and class assignments.

For each training run, multiple model architectures are trained in parallel with different configurations. Our implementation trained six XGBoost-based classifiers with no feature scaling (tree-based models do not benefit from scaling) and varying numbers of selected features: no feature selection (all 1,633 features), top 100 features selected by mutual information, and top 50 features. We also trained linear baselines (logistic regression with StandardScaler) and ensemble models (random forest) for comparison. Feature filtering removes metadata columns (plate, well, tile, cell identifiers, spatial coordinates, and barcode assignments) before training, retaining only morphological measurements. For our mitotic/interphase classifier, features were extracted across all four imaging channels (DAPI, Tubulin, yH2AX, Actin) for nucleus, cell, and cytoplasm compartments, yielding 1,633 features after metadata filtering.

The training pipeline implements stratified 5-fold cross-validation to estimate generalization performance. For each model configuration, the pipeline splits data into training (80%) and test (20%) sets with stratified sampling to preserve class balance, fits the specified scaler on training data if applicable, applies optional feature selection using SelectKBest with mutual information criterion, trains the classifier on transformed training data, evaluates on held-out test data to compute accuracy, precision, recall, and F1 scores for each class, generates confusion matrices and per-class evaluation plots, and serializes the complete pipeline (scaler plus feature selector plus classifier) using dill for deployment. Feature importance analysis is performed for tree-based models, with importance scores visualized as horizontal bar plots showing the top 20 contributing features.

The module optionally supports post-hoc confidence calibration to improve the reliability of predicted probabilities. Two calibration methods are implemented: isotonic regression (recommended for datasets with more than 100 examples) and sigmoid calibration (Platt scaling, suitable for small datasets). Calibration fits a monotonic transformation that maps raw classifier confidence scores to calibrated probabilities that better reflect true classification accuracy. For calibration, a separate labeled dataset (or the training dataset if no calibration set is provided) is used to fit the calibration function. The calibrated classifier is then applied to test data, and calibration quality is assessed by comparing confidence score distributions before and after calibration. In our analysis, we did not apply confidence calibration as the raw XGBoost confidence scores exhibited good agreement with classification accuracy.

Trained classifiers are applied to full datasets through the CellClassifier class, which loads the serialized model pipeline and applies it to feature matrices extracted from phenotype or merge data. The classifier assigns each cell a predicted class label (1-indexed integer corresponding to the class list order) and a confidence score (probability of the predicted class). Per-class confidence thresholds are implemented with two filtering modes: ‘exclude’ mode removes cells with confidence below threshold from downstream analysis, while ‘reassign’ mode tests low-confidence cells against alternative class thresholds and reassigns them if they exceed another class’s threshold, otherwise excluding them. This configuration enables asymmetric filtering strategies where minority classes such as mitotic cells are held to stricter standards while majority classes such as interphase cells use more permissive thresholds to maximize cell retention.

The rankline UI provides an interactive tool for empirically determining appropriate confidence thresholds. Cells are displayed in a scrollable grid sorted by prediction confidence (descending order for each class), with actual images shown alongside predicted class and confidence score. Users can visually inspect high-confidence and low-confidence predictions to identify the confidence level at which classification errors become frequent, then set thresholds accordingly. The UI supports filtering by class, minimum confidence difference to focus on ambiguous cases, and direct navigation to specific confidence ranges.

Classified cells are automatically partitioned by the Aggregate module based on their assigned class labels. Each class is processed independently through the aggregation pipeline, generating separate gene-level embeddings for each cell state. This enables cell-state-specific phenotypic analyses, such as separate clustering of interphase and mitotic cell populations to identify genes with distinct perturbation effects in different cell cycle stages. For our analysis, we configured the classifier to create two subpopulations (mitotic and interphase), which were independently aggregated and clustered using the parameters optimized for each cell state.

### Aggregate Module

The Aggregate module transforms single-cell perturbation data into robust gene-level embeddings through a systematic workflow that enables analysis of specific cellular subpopulations. It accepts classified cell populations from the Classify module and can further subdivide datasets by channel combinations of interest, enabling analysis of phenotypic effects specific to particular cellular structures or markers.

The filter step implements a comprehensive strategy addressing common data quality issues. Query-based filtering allows removal of specific cell subsets based on configurable criteria; our implementation filtered to cells with a single gene mapping. Perturbation filtering restricts analysis to cells with assigned genetic perturbations, while missing value filtering drops columns and rows with a configurable proportion of missing data (we used thresholds of 0.05 for columns and 0.01 for rows). The module offers optional imputation of remaining missing values using K-nearest neighbors with k=5, implemented in batches (default 1000 rows per batch with 10000 sampled complete cases) to handle large datasets efficiently (enabled in our implementation). The Local Outlier Factor algorithm identifies outlier cells with anomalous marker patterns for removal, using mean channel intensities as input features, with a configurable contamination parameter (set to 0.01).

The module optionally implements per-cell perturbation scoring to assess the strength of each cell’s perturbation phenotype, which is particularly valuable for perturbations with heterogeneous penetrance or for identifying off-target effects. For each gene, the algorithm samples an equal number of non-targeting control cells matched to the perturbed cell count, centers and scales features on controls within each plate-well batch, selects the top 200 differential features using ANOVA F-test, trains a logistic regression classifier (L2 regularization, balanced class weights, max 2000 iterations) to distinguish perturbed from control cells, generates out-of-fold probability predictions via stratified k-fold cross-validation, and computes the ROC AUC as a measure of overall perturbation strength. The resulting per-cell perturbation scores (probability of being perturbed) and gene-level AUC metrics enable fine-grained quality control. Cells with perturbation scores below a specified threshold can be filtered before aggregation, removing cells with weak or noisy phenotypic effects that would dilute gene-level profiles. Genes with insufficient cell counts (less than 100 cells by default) receive NaN scores and are retained without filtering. The perturbation scoring implementation processes genes in parallel batches to manage memory usage, with each batch independently computing scores before updating the main dataset. For our analysis, we did not apply perturbation score filtering so as to more tightly compare to the previous analysis, retaining all cells with assigned gene mappings. However, when conducting larger genome-wide screens, this becomes a tractable way to subset to perturbations that result in a real morphological deviation from control cells.

Batch alignment addresses technical variation by first organizing samples by experimental batch (defined in our case by plate and well identifiers), then applying Principal Component Analysis (PCA) to reduce dimensionality while preserving variation. For large datasets exceeding 50,000 cells, PCA is first fit on a subsample of 50,000 cells to determine the number of components, then the full dataset is transformed. Users can configure either the target variance to preserve (we used 0.99) or a specific component count.

The critical Typical Variation Normalization (TVN) step leverages non-targeting control samples to standardize data through a multi-step process: (1) center and scale all data based on control sample mean and standard deviation; (2) fit PCA on control samples and transform all data to this space; (3) center and scale again based on controls (optionally per batch); (4) for each batch, compute the source covariance matrix from batch controls, then apply the transformation 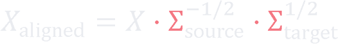, where Σ_target_ is the global control covariance. The matrix fractional powers are computed using scipy’s fractional_matrix_power function. A small regularization term (0.5 × identity matrix) is added to covariance matrices to ensure numerical stability. This generates globally representative principal components at the single-cell level that capture essential morphological feature relationships while removing technical variation.

The final aggregation stage combines cellular measurements into perturbation-level representations through two complementary approaches. First, batch-normalized principal components are aggregated to the perturbation level to create embeddings used for clustering analysis that reveals functional relationships between genetic manipulations. Second, the original interpretable features (such as nuclear size and stain intensities) are aggregated to the perturbation level to provide quantitative metrics that directly inform understanding of which specific cellular processes drive observed functional clusters.

The aggregation supports multiple statistical methods to accommodate different analytical objectives: mean aggregation provides the arithmetic average of cellular measurements, appropriate when the goal is to capture the central tendency of normally distributed features; median aggregation offers robustness to outliers and skewed distributions, which we selected for our analysis; and trimmed mean aggregation excludes extreme values before averaging, providing a balance between sensitivity and robustness. For experiments with variable cell counts per perturbation, weighted aggregation schemes can incorporate cell count as a weighting factor.

To assess the stability and significance of aggregated features, the module implements bootstrapping at the feature level. For each perturbation, cells are resampled with replacement across multiple iterations, and aggregation statistics are computed for each resample. This generates empirical distributions for each aggregated feature, from which confidence intervals and standard errors can be derived. The bootstrapped distributions enable statistical comparison between perturbations, identification of features with high measurement uncertainty, and assessment of whether observed differences exceed sampling variability. Bootstrap-derived p-values can be computed by comparing each perturbation’s feature distribution against the non-targeting control distribution, with multiple testing correction applied across features and perturbations.

Quality assessment tools are integrated throughout the module, generating comprehensive reports of missing value patterns and feature distributions before and after batch correction. The implementation leverages efficient data structures and lazy evaluation to handle large datasets, with support for parallel processing and memory optimization for high-performance computing environments.

### Cluster Module

The Cluster module identifies functional relationships between genes by grouping genetic perturbations with similar phenotypic profiles. This final analytical stage transforms the high-dimensional embeddings from the Aggregate module into interpretable clusters that reveal biological pathways and functional connections.

At the core of the Cluster module is a dimensionality reduction and clustering pipeline built on PHATE (Potential of Heat-diffusion for Affinity-based Transition Embedding) and Leiden community detection. PHATE is run with k=10 nearest neighbors and a configurable distance metric, producing both a 2D embedding and a diffusion potential matrix that captures the underlying manifold structure. PHATE preserves both local and global structure while reducing high-dimensional data to a visually interpretable two-dimensional representation. This approach leverages a diffusion-based metric that captures continuous phenotypic transitions better than linear methods like PCA or nonlinear methods like t-SNE.

The PHATE graph weights serve as input to the Leiden algorithm. Before clustering, the weight matrix is symmetrized by averaging with its transpose: *W_symmetric_* = (*W* + *W^T^*)/2. This symmetric matrix is then converted to an undirected weighted graph, and the Leiden algorithm is applied using the RBConfigurationVertexPartition method with a configurable resolution parameter controlling cluster granularity. The implementation uses the leidenalg library with n_iterations=-1 for convergence.

To objectively evaluate clustering quality, the module implements a comprehensive benchmarking framework that tests gene clusters against established biological databases. The evaluation employs two complementary approaches: pairwise gene relationship analysis using STRING protein-protein interactions, and group-level enrichment analysis using CORUM protein complexes and KEGG pathways. For pairwise evaluation, the system identifies true positive interactions when genes from known interacting pairs appear in the same cluster, while false positives and negatives are determined by mismatched clustering relative to known interactions. This yields standard precision and recall metrics, with an optional adjusted precision calculation that accounts for incomplete knowledge in reference databases. For group-level evaluation, the module applies Fisher’s exact test with Benjamini-Hochberg correction to assess enrichment of known gene complexes or pathways within each cluster. The system filters reference complexes to ensure robustness, requiring at least three genes present in the dataset and at least two-thirds of complex components represented. An additional filter removes larger complexes that share more than 10% of gene pairs with smaller complexes, preventing redundant enrichment calls. The pipeline performs parallel analysis against randomly shuffled controls to establish baseline performance and confirm that identified patterns represent true biological signal rather than technical artifacts.

Beyond clustering, the module computes a phenotypic distance metric for each perturbation relative to non-targeting controls using the PHATE diffusion potential space. For each gene, the average Euclidean distance from its potential vector to all non-targeting control potential vectors is calculated, providing a quantitative measure of phenotypic divergence. These distances are then min-max normalized across all perturbations to yield a 0-1 scaled “potential distance” score, where higher values indicate stronger phenotypic effects. This metric serves as a continuous measure of perturbation strength that complements discrete cluster assignments.

To focus analyses on perturbations with significant phenotypic effects, the module implements filtering based on the per-gene AUC scores computed during the Aggregate module’s perturbation scoring step. Perturbations with AUC values below a user-specified threshold (e.g., 0.6 or 0.7) are excluded from downstream clustering and visualization, restricting analysis to genes that demonstrate distinguishable phenotypes relative to non-targeting controls. This filtering is particularly valuable for large screens where all cells are retained through aggregation without perturbation score-based cell filtering; by removing low-AUC perturbations before clustering, researchers can reduce noise from inactive or poorly penetrant knockouts and focus on genes most likely to yield interpretable biological insights.

The Cluster module provides extensive visualization capabilities, generating PHATE embeddings that allow intuitive exploration of gene relationships in two-dimensional space. Cluster assignments are highlighted through color coding, enabling rapid identification of gene communities. When AUC filtering is applied, filtered embeddings display only perturbations exceeding the significance threshold, with non-targeting controls shown for reference. Additional visualizations include cluster size distribution plots to assess clustering granularity, enrichment pie charts showing the proportion of clusters supported by different reference databases, precision-recall curves to identify optimal clustering resolutions, and AUC distribution histograms to characterize the overall phenotypic effect landscape of the screen. These visualization tools facilitate both quality control assessment and biological interpretation of the clustering results.

### MozzareLLM

As part of the Brieflow pipeline, we implemented MozzareLLM, a framework that leverages large language models to analyze phenotypic clusters and prioritize gene candidates for experimental validation. MozzareLLM employs a structured prompt engineering approach organized into sequential analytical stages.

For each cluster, the model is provided with the list of member genes along with their UniProt functional annotations, which supply the primary biological context for reasoning about gene function and pathway membership. The model is first tasked with identifying the dominant biological pathway that explains why the genes in the cluster exhibit similar phenotypic profiles. Pathway confidence is assessed according to stringent criteria based on the proportion of genes that fit the proposed pathway: high confidence requires greater than 70% of genes with strong literature support, medium confidence requires 50–70%, and low confidence requires 30–50%, with clusters below 30% designated as having no coherent pathway.

The model is then tasked with classifying each gene into one of three mutually exclusive categories relative to the identified pathway. Established pathway genes are well-documented members with clear functional roles supported by multiple publications. Uncharacterized genes have minimal to no functional annotation, with limited experimental validation and few publications. Novel role genes have established functions in other pathways but may contribute to the identified pathway in a previously unrecognized way. Classifications are informed by the UniProt annotations provided in the prompt, which the model uses to assess the extent of existing functional characterization for each gene.

For both uncharacterized and novel role genes, the model assigns a prioritization score from 1 to 10. Uncharacterized genes receive higher scores when they are virtually unstudied with unknown molecular function, and lower scores when partial evidence for pathway involvement exists. Novel role genes receive higher scores when a compelling rationale exists for a previously unrecognized role with minimal existing literature, and lower scores when existing data already supports involvement in the pathway.

To select the optimal language model for this task, we evaluated three frontier large language models—Claude Opus 4.6 (Anthropic), GPT-5.2 (OpenAI), and Gemini 3 Pro Preview (Google)—on their ability to annotate gene clusters from functional genomics screens. Each model was tasked with identifying the dominant biological process for clusters drawn from three benchmark datasets: Optical Pooled Screens (OPS; 7 clusters)^9^, DepMap genetic dependencies (2 clusters)^31^, and proteomics co-complex data (2 clusters)^32^, totaling 11 ground-truth-annotated and published clusters. Claude Opus 4.6 and GPT-5.2 performed equivalently, each correctly identifying the dominant biological process for 10 of 11 clusters (91%), while Gemini 3 Pro Preview matched 9 of 11 (82%). All three models appropriately identified the dominant biological processes in the DepMap and Proteomics benchmarks. Performance differences emerged exclusively in the OPS benchmark, which contains more heterogeneous clusters spanning multiple sub-complexes. Gemini failed to identify the mTOR signaling pathway in a multi-complex cluster (cluster 37), returning “Unknown,” whereas both Claude Opus and GPT-5.2 correctly synthesized the mTOR/Golgi trafficking/Integrator sub-complex architecture. The sole cluster missed by all three models (cluster 197) had a ground-truth annotation of m6A mRNA modification—all models instead identified cell cycle and chromatin regulation as the dominant process, reflecting the greater representation of these genes within the cluster. Based on equivalent performance and platform stability, we selected Claude Opus 4.6 (temperature=0.0) for all cluster annotations in the current analysis.

Our implementation supports integration with multiple large language model providers including OpenAI, Anthropic, and Google, allowing researchers to select the most appropriate model for their analytical needs. The MozzareLLM methodology produces standardized outputs that include the dominant biological process, pathway confidence level, gene classifications with prioritization scores, and rationales explaining why particular genes merit investigation. This structured approach facilitates efficient experimental planning by highlighting the most promising candidates for functional validation, particularly those representing potential novel pathway components or genes with unexpected functions.

To verify that high-confidence pathway annotations reflect genuine biological signal rather than spurious pattern recognition, we performed a negative control experiment. We shuffled cluster assignments while preserving cluster size distributions, destroying biological relationships while maintaining the overall structure of the embedding space. MozzareLLM was then applied to these scrambled clusters using identical parameters (Claude Opus 4.6, temperature=0.0). This scrambled validation provides objective evidence that the pathway interpretations require genuine functional relationships rather than arising from random gene associations or LLM hallucination. For each benchmark dataset, the screen context provided to the model was adapted to reflect the specific experimental design of that dataset. In evaluation of the clusters generated by Brieflow, the prompt additionally included the imaging markers used and the cell population (interphase or mitotic) for each cluster. All comparisons between Funk et al. clusters and Brieflow clusters were performed using identical prompts.

### HeLa Screen Generation and Analysis

The ‘Vesuvius’ HeLa screen dataset comprised a CRISPR-Cas9 screen targeting 5,072 fitness-conferring genes with four sgRNAs per gene and 250 non-targeting control sgRNAs. Following library transduction into HeLa cells with integrated doxycycline-inducible Cas9, cells underwent puromycin selection for 4 days. Cas9 expression was then induced with doxycycline for 78 hours, identified as the optimal timepoint for maximizing observable phenotypes while minimizing dropout of knockout cells. The dataset consisted of eight 6-well plates with 46 wells containing fixed-cell imaging data capturing four cellular markers (DNA, DNA damage response, actin, and microtubules). More comprehensive details on how the screen was conducted are available in the original publication^9^.

---

**Extended Data Fig. 1:**
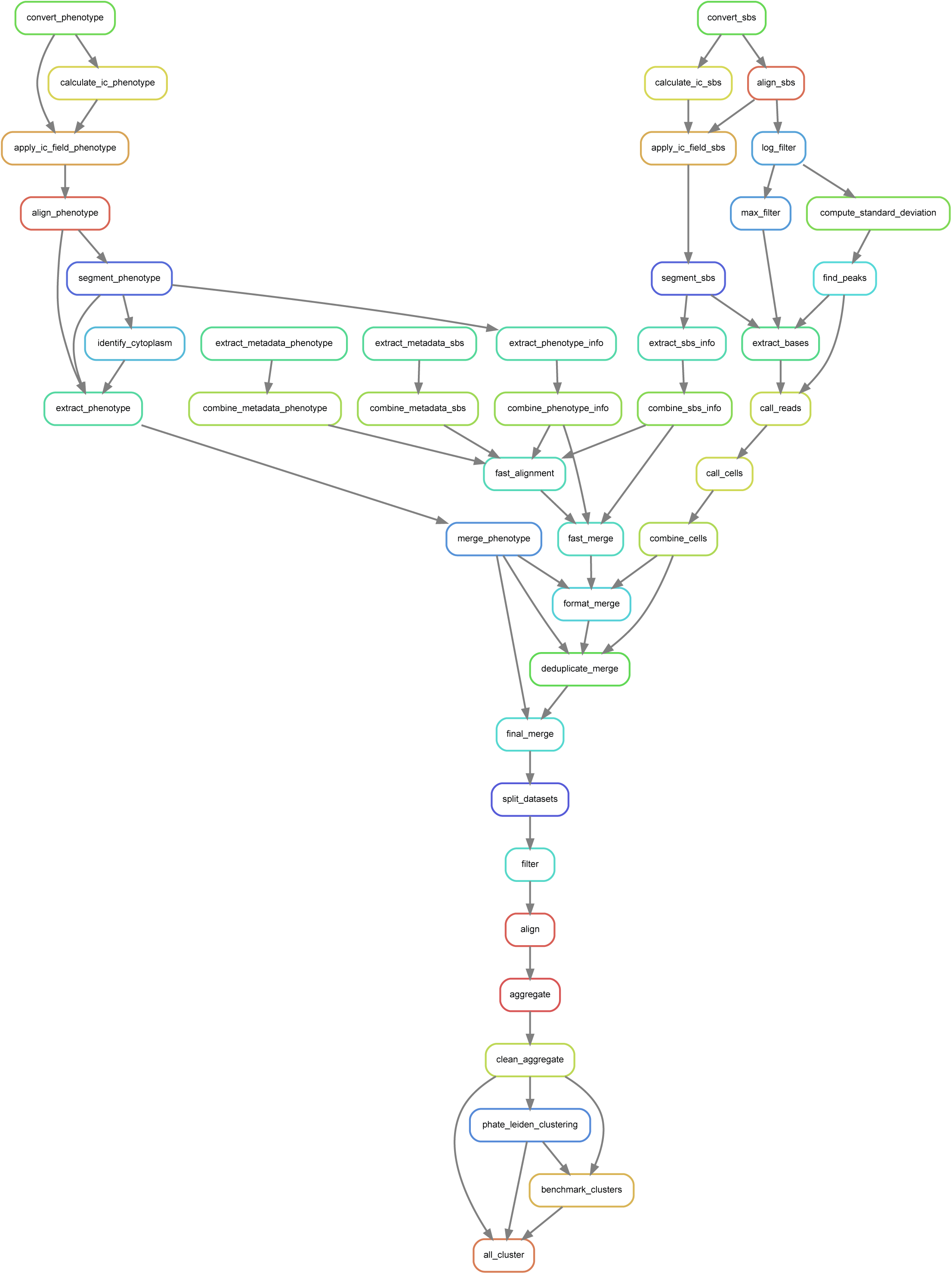
Snakemake-generated rule graph for the Brieflow analysis pipeline, showing the directed acyclic graph of all processing steps from raw data conversion through final clustering and benchmarking.

**Extended Data Fig. 2:**
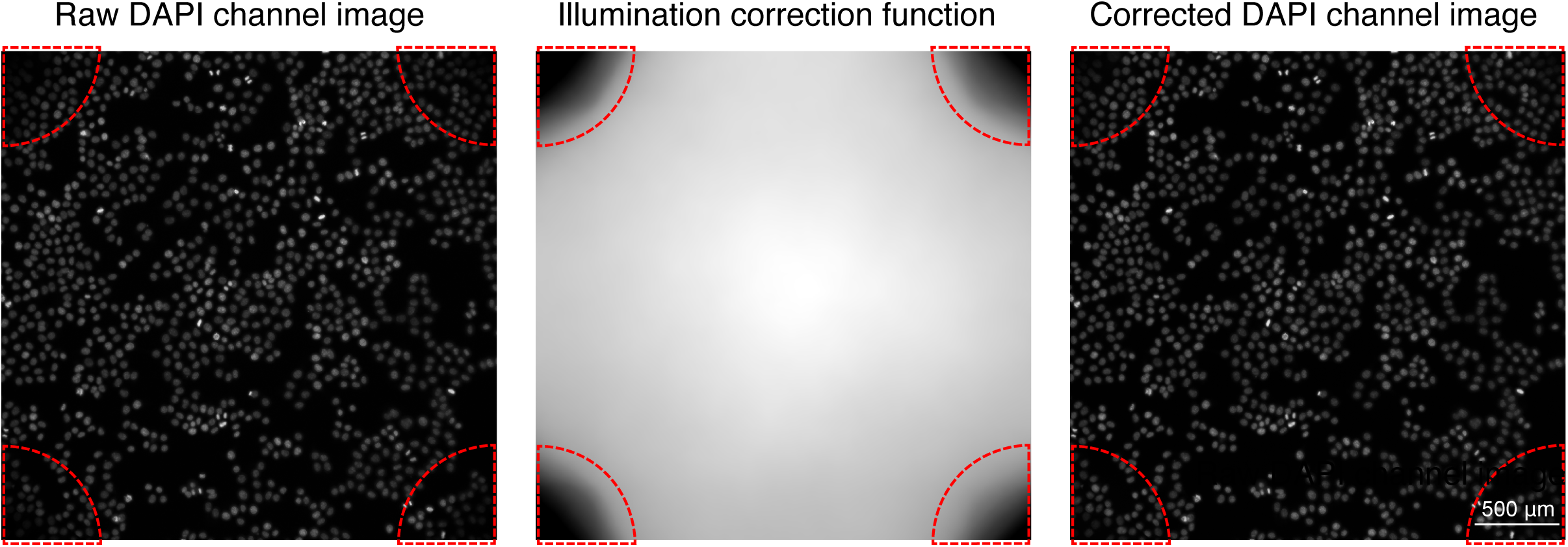
Illumination correction for phenotypic image processing. Raw DAPI channel image (left), the computed illumination correction function (center), and the corrected DAPI channel image (right) for a representative field of view. Red dashed circles highlight well corners where illumination intensity is reduced, demonstrating the correction of vignetting artifacts.

**Extended Data Fig. 3:**
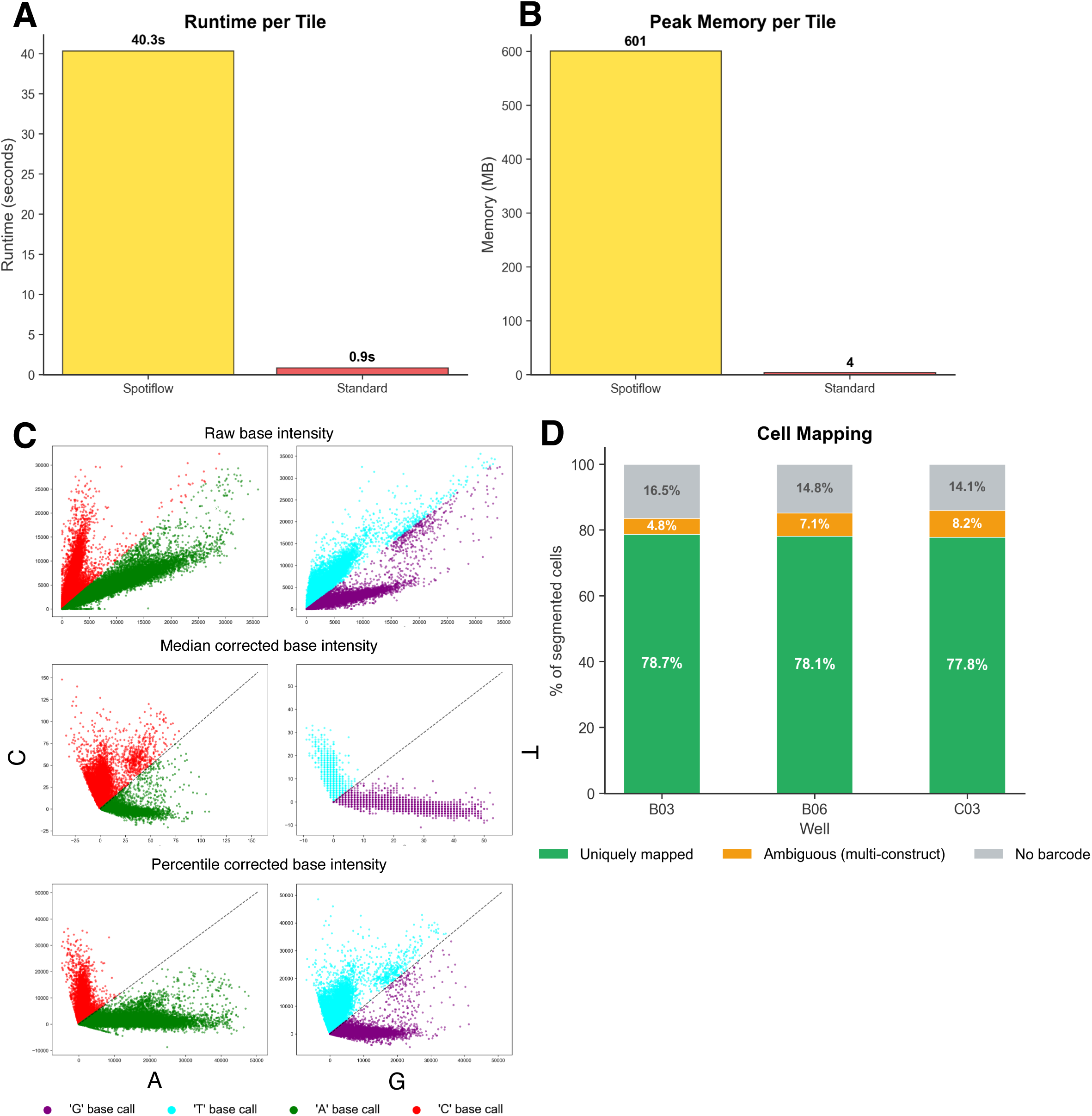
Benchmarking of spot detection methods and normalization strategies. **a.** Runtime comparison between Spotiflow and the standard method, showing substantially longer processing time for Spotiflow (40.3 s vs. 0.9 s per tile). **b.** Peak memory usage comparison, showing higher memory consumption for Spotiflow (601 MB vs. 4 MB per tile). **c.** Comparison of normalization strategies for base calling. Raw fluorescence intensities (top row) are transformed using median normalization (middle row) or percentile normalization (bottom row). Left panels show intensity distributions for bases A and C; right panels show bases T and G. Both approaches improve discrimination between complementary base pairs, with colored points indicating base assignments. **d.** Validation of the Sequencing-by-Synthesis module on an independent T7-based in situ sequencing dataset with three known barcode constructs. Stacked bar plots show the proportion of segmented cells uniquely mapped to a single construct, ambiguously mapped to multiple constructs, or unmapped, across three wells.

**Extended Data Fig. 4:**
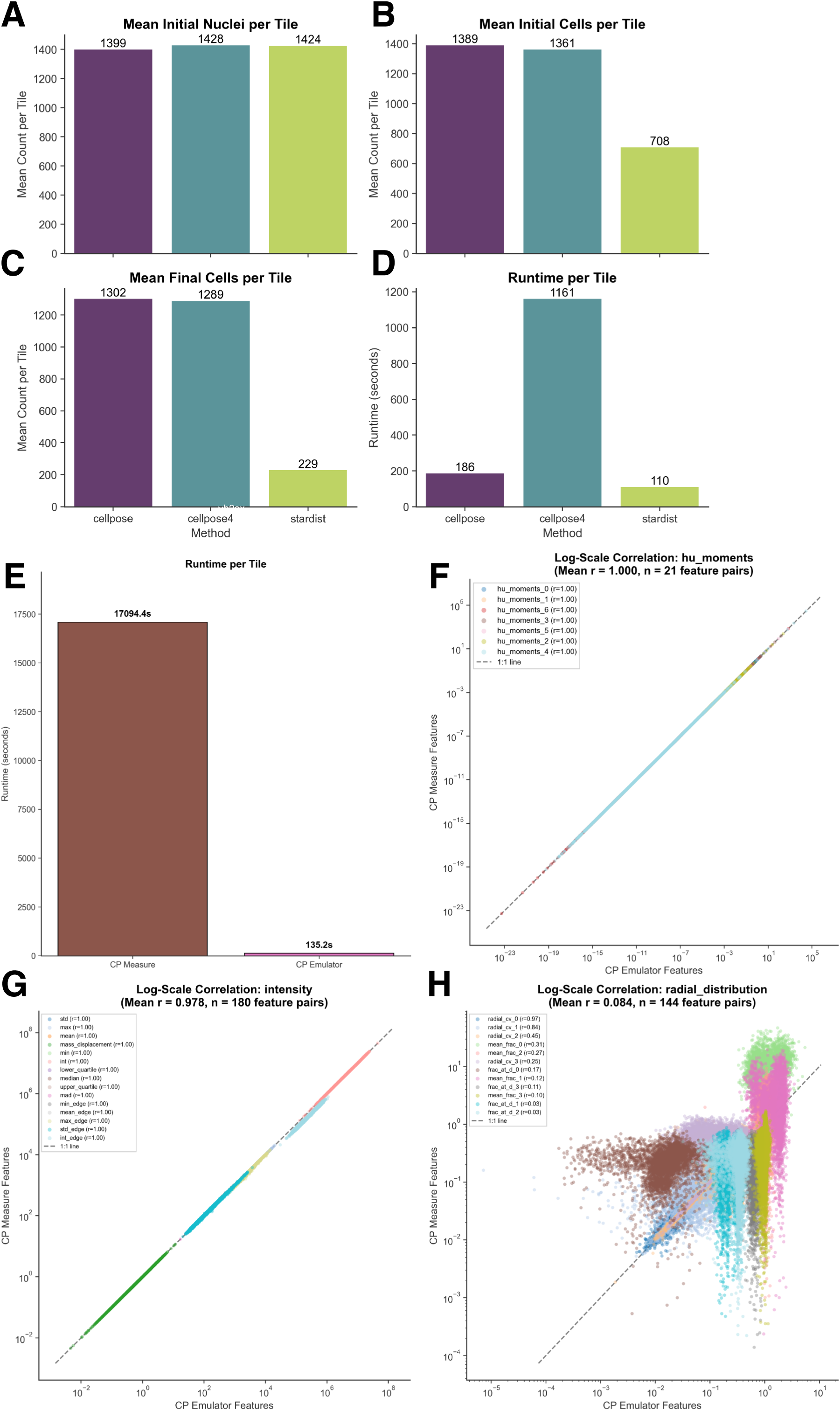
Comparative analysis of segmentation methods and feature extraction implementations. **a.** Mean initial nuclei detected per tile across Cellpose, Cellpose4, and StarDist, showing similar counts. **b.** Mean initial cells detected per tile, showing substantially fewer cells detected by StarDist. **c.** Mean final cells per tile after reconciliation of cell and nuclear measurements, showing that Cellpose and Cellpose4 retain most cell–nuclei pairs while StarDist shows a large reduction. **d.** Runtime comparison of segmentation methods, with Cellpose4 substantially slower than Cellpose and StarDist. **e.** Runtime comparison between cp_measure and the Brieflow emulator for feature extraction, showing approximately 125-fold faster processing by the emulator. **f.** Log-scale correlation of cellular Hu moment features between the Brieflow emulator and cp_measure, showing strong agreement (mean r = 1.000). **g.** Log-scale correlation of cellular intensity features between implementations, showing high agreement (mean r = 0.978). **h.** Log-scale correlation of cellular radial distribution features between implementations, showing minimal agreement (mean r = 0.084). Each dot in **f–h** represents one cell from 8 benchmark tiles, with colors indicating distinct feature measurements and r denoting the Pearson correlation coefficient.

**Extended Data Fig. 5:**
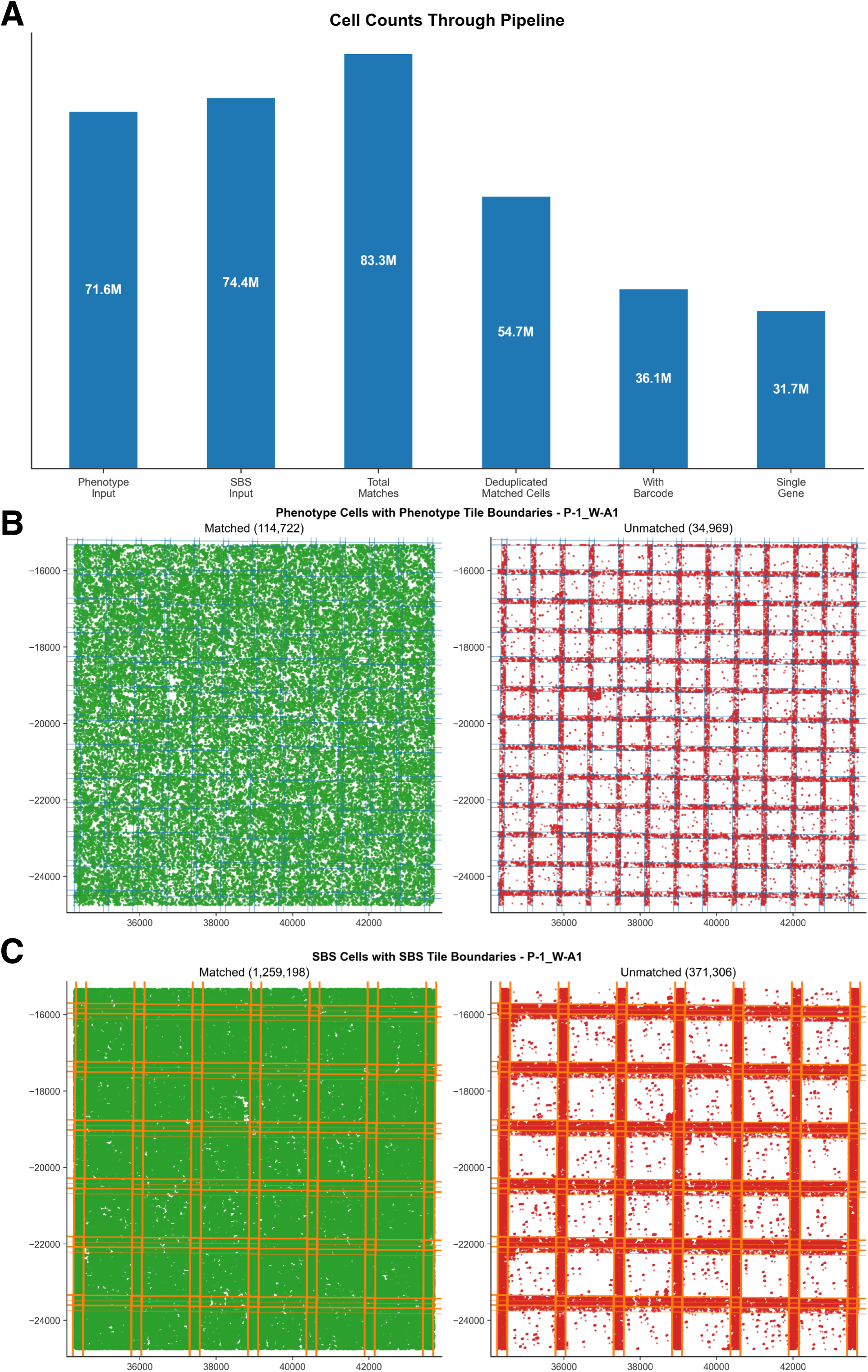
Merge performance and spatial registration validation. **a.** Cell counts at each stage of the Brieflow pipeline, from Phenotype input (71.6M) and SBS input (74.4M) through total matches (83.3M), deduplicated matched cells (54.7M), cells with barcode (36.1M), to cells with single gene assignment (31.7M). **b.** Spatial distribution of matched (left, green) and unmatched (right, red) phenotype cells for a representative well, with phenotype tile boundaries overlaid. Unmatched cells concentrate at tile boundaries where overlapping fields of view produce duplicate detections. **c.** Spatial distribution of matched (left, green) and unmatched (right, red) SBS cells for the same well, with SBS tile boundaries overlaid, confirming that cell loss during merge reflects boundary deduplication rather than registration failure.

**Extended Data Fig. 6:**
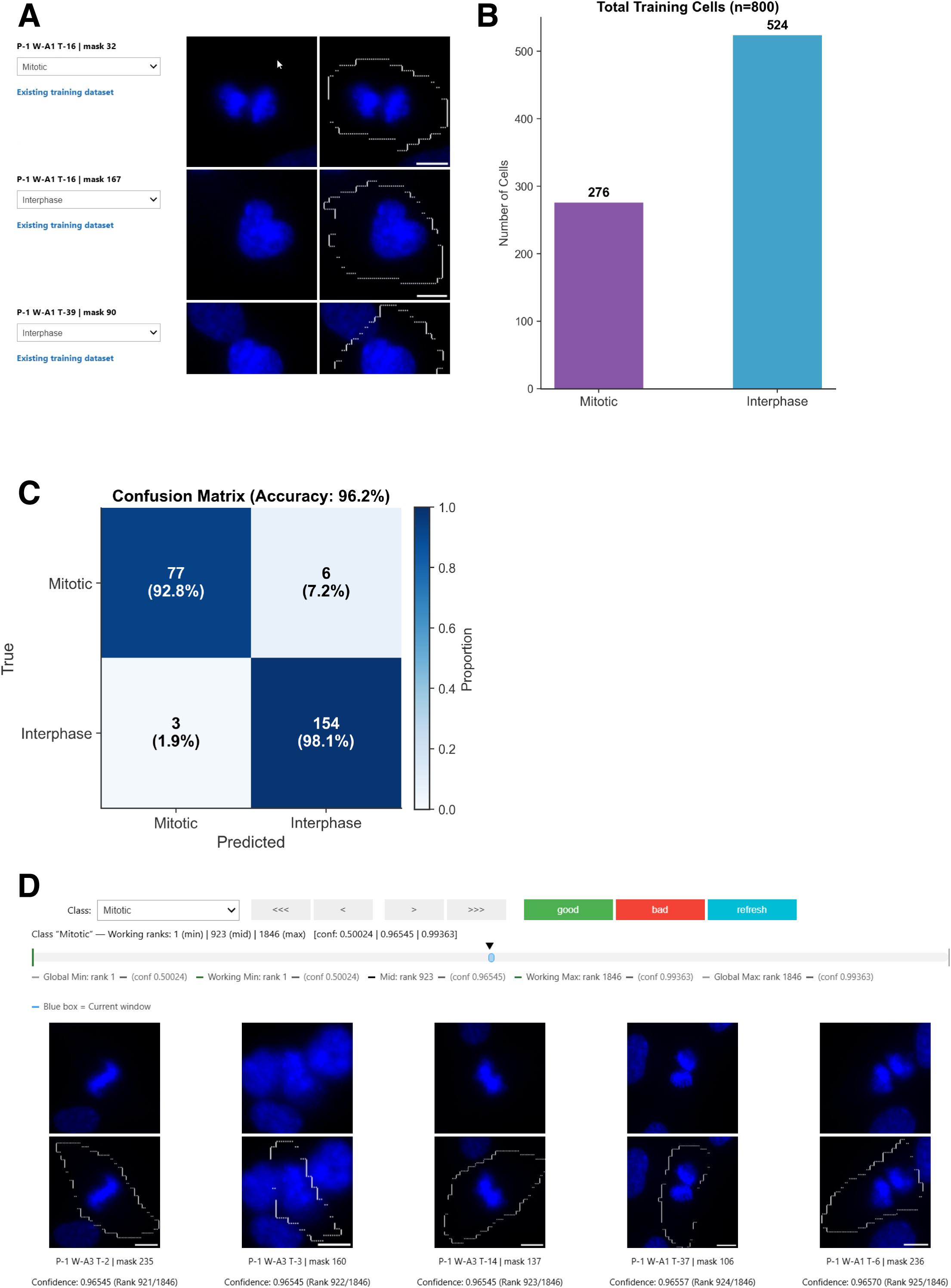
Development and validation of a mitotic cell classifier. **a.** Interactive labeling interface showing representative cell images across multiple channels, with class assignment dropdown and existing dataset indicators. **b.** Training dataset composition (n = 800 cells total: 276 mitotic, 524 interphase). **c.** Confusion matrix on the held-out test set, showing 96.2% overall accuracy with 92.8% sensitivity for mitotic cells and 98.1% sensitivity for interphase cells. **d.** Rankline interface for empirical confidence threshold determination, displaying mitotic cells ranked by prediction confidence with cell images across DAPI and tubulin channels.

**Extended Data Fig. 7:**
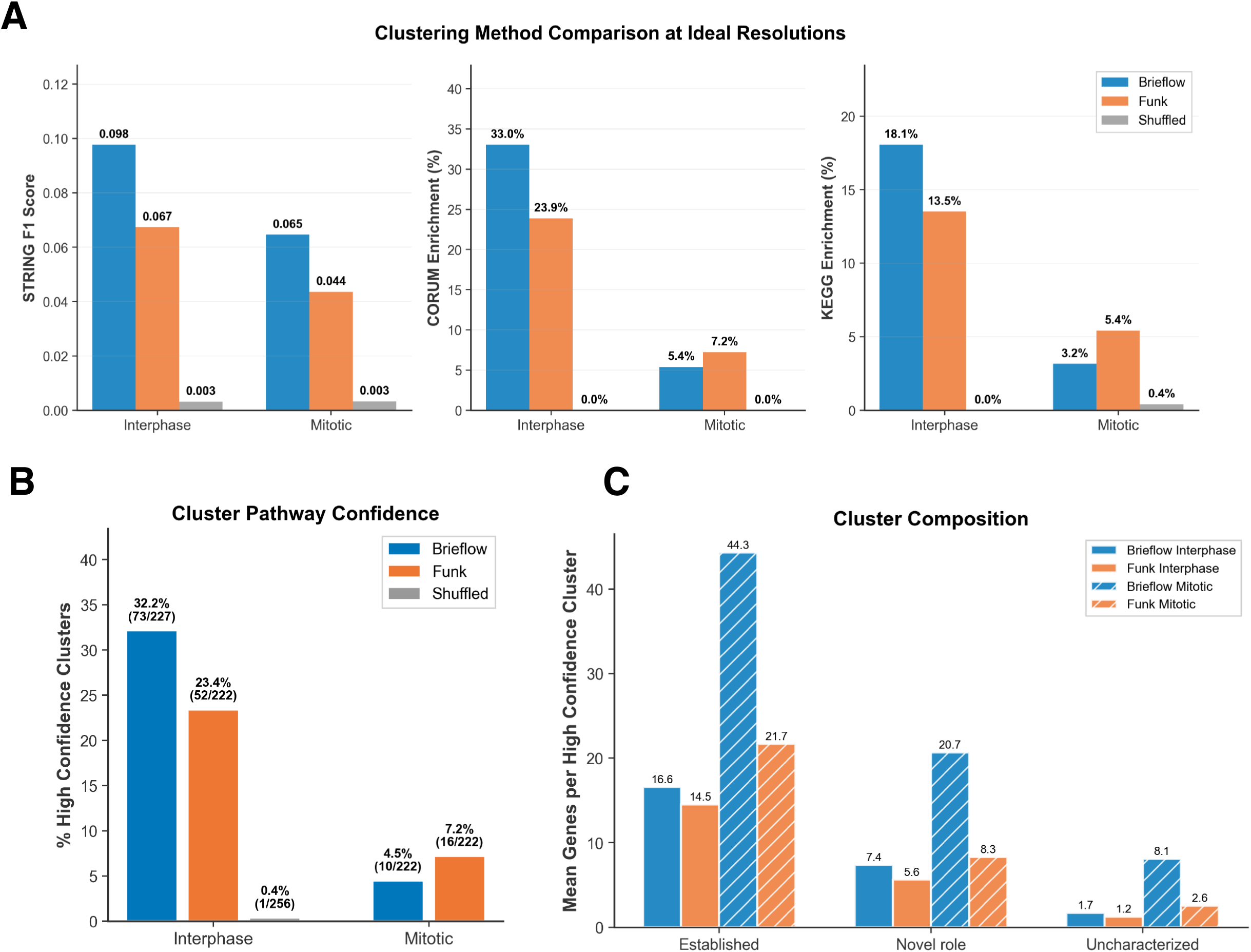
Clustering benchmarks, MozzareLLM validation, and gene classification. **a.** Comparison of clustering quality metrics between Brieflow, Funk et al., and shuffled controls at their respective optimal resolutions. Left: STRING F1 scores for interphase and mitotic populations. Center: CORUM enrichment. Right: KEGG enrichment. Shuffled controls show near-zero scores across all metrics. **b.** Percentage of clusters receiving high-confidence MozzareLLM pathway annotations for Brieflow interphase (32.2%), Funk et al. interphase (23.4%), shuffled interphase (0.4%), Brieflow mitotic (4.5%), and Funk et al. mitotic (7.2%). **c.** Mean number of genes per high-confidence cluster classified as established, novel-role, or uncharacterized, shown for Brieflow and Funk et al. across interphase and mitotic populations.

**Extended Data Fig. 8:**
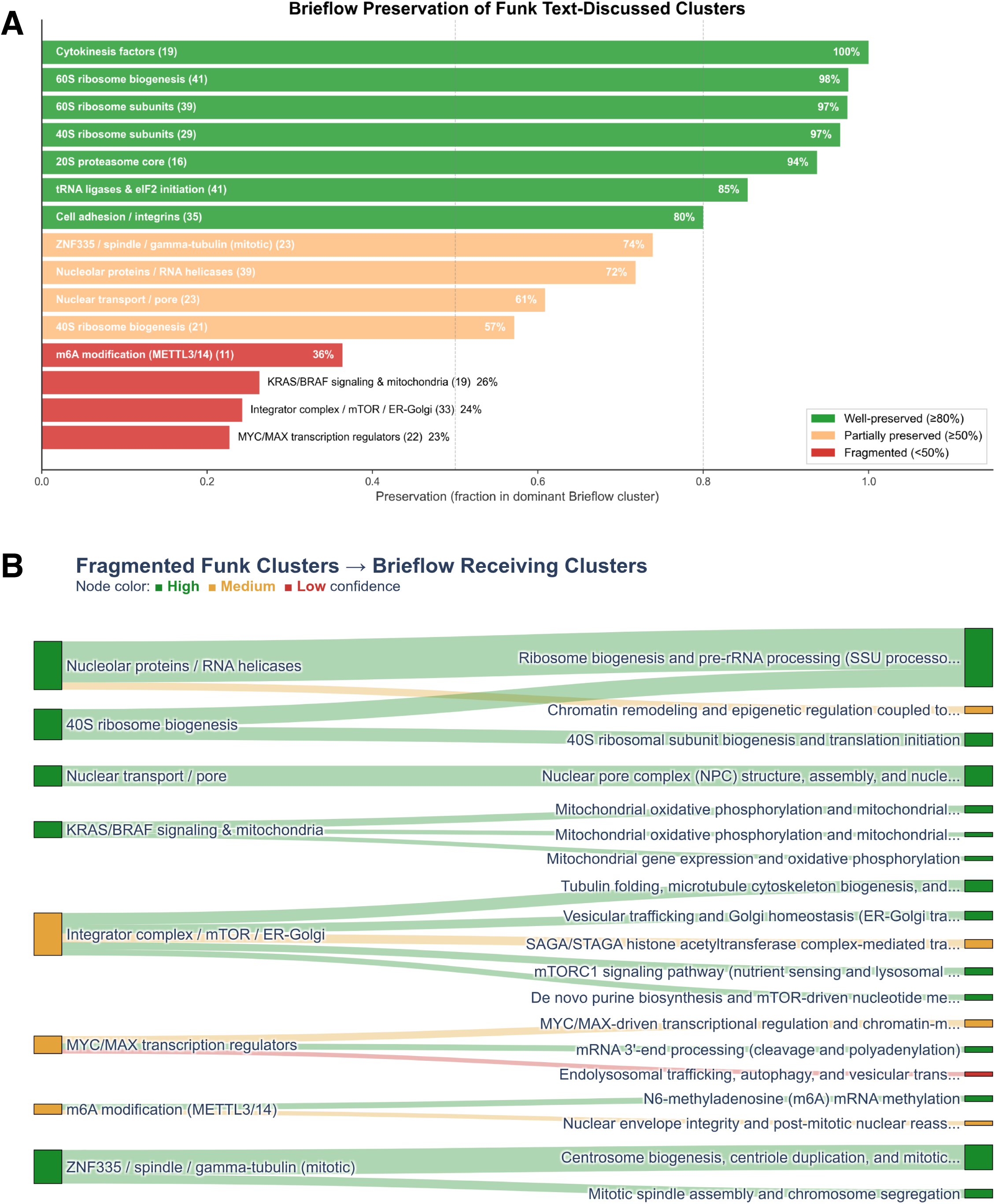
Preservation of Funk et al. biological findings in Brieflow clustering. **a.** Preservation analysis of the 15 Funk et al. text-discussed clusters, showing the fraction of each cluster’s genes that co-occur in a single dominant Brieflow cluster. Clusters are colored by preservation level: well-preserved (_≥_80%, green), partially preserved (_≥_50%, yellow), or fragmented (<50%, red). **b.** Redistribution of genes from fragmented Funk et al. clusters into Brieflow clusters. Left nodes represent original Funk et al. clusters; right nodes represent receiving Brieflow clusters, colored by MozzareLLM pathway confidence (high, medium, or low). The majority of redistributed genes are reassigned to high-confidence Brieflow modules.

**Extended Data Fig. 9:**
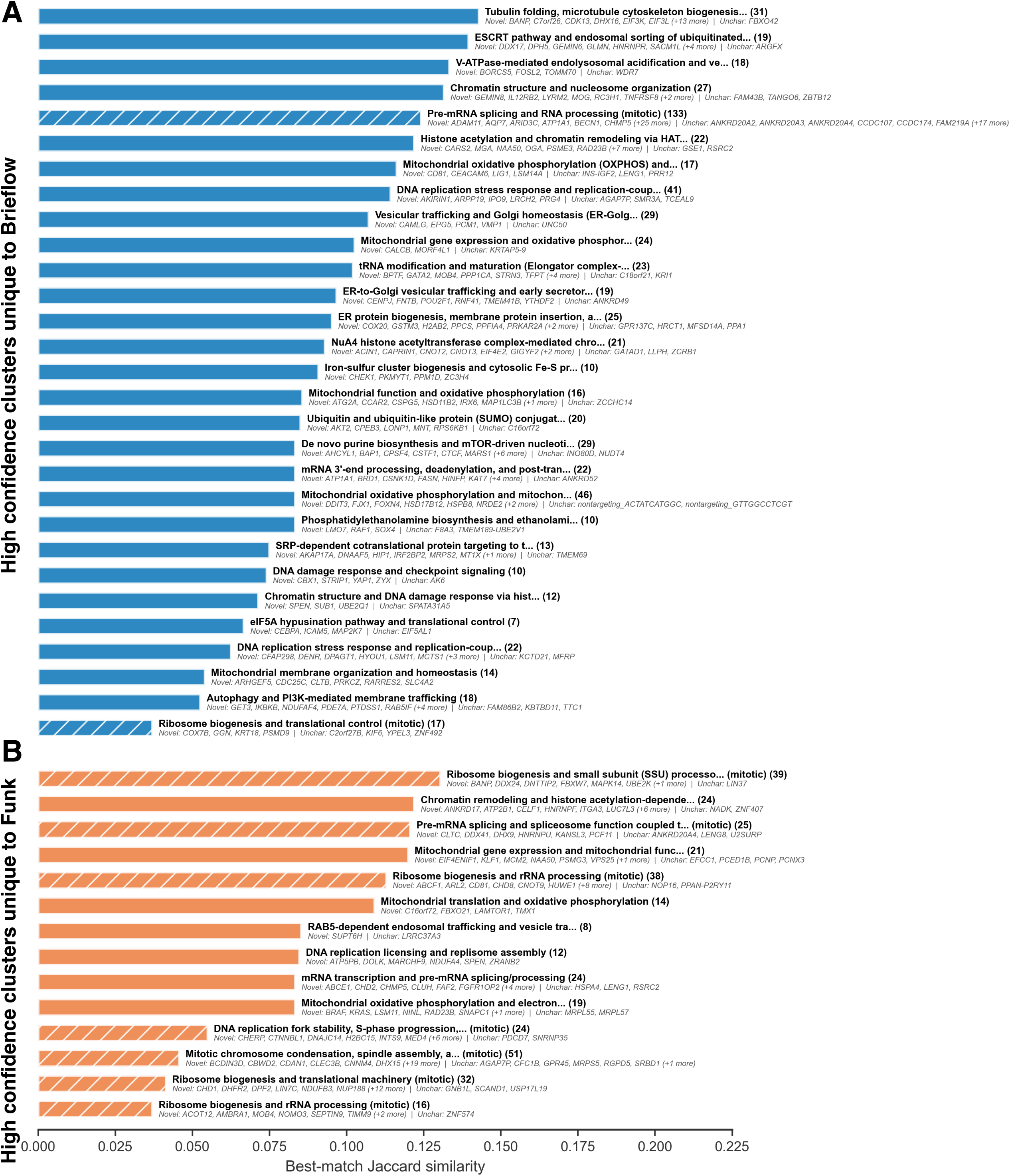
Novel functional programs resolved by Brieflow. **a.** High-confidence Brieflow clusters with no adequate match (Jaccard < 0.15) in the Funk et al. clustering, ranked by best-match Jaccard similarity. Each bar is annotated with the cluster pathway name, size, and lists of novel-role and uncharacterized genes. **b.** High-confidence Funk et al. clusters with no adequate match in the Brieflow clustering, shown in the same format. The asymmetry (29 Brieflow-unique vs. 14 Funk-unique clusters) reflects Brieflow’s finer functional resolution.

## References

1. Feldman, D. et al. Optical Pooled Screens in Human Cells. Cell 179, 787–799.e17 (2019).

2. Carlson, R. J., Leiken, M. D., Guna, A., Hacohen, N. & Blainey, P. C. A Genome-Wide Optical Pooled Screen Reveals Regulators of Cellular Antiviral Responses. Proceedings of the National Academy of Sciences of the United States of America 120, 2210623120 (2023).

3. Ramezani, M. et al. A genome-wide atlas of human cell morphology. Nat Methods 22, 621–633 (2025).

4. Labitigan, R. L. D. et al. Mapping variation in the morphological landscape of human cells with optical pooled CRISPRi screening. eLife 13, (2024).

5. Wang, Z. J. et al. Multi-ContrastiveVAE disentangles perturbation effects in single cell images from optical pooled screens. 2023.11.28.569094 Preprint at 10.1101/2023.11.28.569094 (2024).

6. Bigverdi, M., et al. Gene-Level Representation Learning via Interventional Style Transfer in Optical Pooled Screening. Preprint at 10.48550/arXiv.2406.07763 (2024).

7. Sivanandan, S. et al. A pooled Cell Painting CRISPR screening platform enables de novo inference of gene function by self-supervised deep learning. Nat Commun 17, 77 (2025).

8. Bunne, C. et al. How to build the virtual cell with artificial intelligence: Priorities and opportunities. Cell 187, 7045–7063 (2024).

9. Funk, L. et al. The phenotypic landscape of essential human genes. Cell 185, 4634–4653.e22 (2022).

10. Mölder, F., et al. Sustainable Data Analysis with Snakemake. (2021).

11. Singh, S., Bray, M.-A., Jones, T. R. & Carpenter, A. E. Pipeline for Illumination Correction of Images for High-Throughput Microscopy. Journal of Microscopy 256, 231–36 (2014).

12. Mantes, A. D. et al. Spotiflow: accurate and efficient spot detection for fluorescence microscopy with deep stereographic flow regression. 2024.02.01.578426 Preprint at 10.1101/2024.02.01.578426 (2024).

13. Kudo, T. et al. Multiplexed, image-based pooled screens in primary cells and tissues with PerturbView. Nat Biotechnol 43, 1091–1100 (2025).

14. Stringer, C. & Pachitariu, M. Cellpose3: One-Click Image Restoration for Improved Cellular Segmentation. Nature Methods 22, 592–99 (2025).

15. Cellpose-SAM: superhuman generalization for cellular segmentation | bioRxiv. https://www.biorxiv.org/content/10.1101/2025.04.28.651001v1.

16. Weigert, M. & Schmidt, U. Nuclei instance segmentation and classification in histopathology images with StarDist. in 2022 IEEE International Symposium on Biomedical Imaging Challenges (ISBIC) 1–4 (2022). doi:10.1109/ISBIC56247.2022.9854534.

17. Stirling, D. R. et al. CellProfiler 4: improvements in speed, utility and usability. BMC Bioinformatics 22, 433 (2021).

18. Muñoz, A. F., et al. cp_measure: API-first feature extraction for image-based profiling workflows. Preprint at 10.48550/arXiv.2507.01163 (2025).

19. Celik, S. et al. Building, Benchmarking, and Exploring Perturbative Maps of Transcriptional and Morphological Data. PLoS Computational Biology 20, 1012463 (2024).

20. Song, B. et al. Decoding heterogeneous single-cell perturbation responses. Nat Cell Biol 27, 493–504 (2025).

21. Ando, D. M., McLean, C. Y. & Berndl, M. Improving Phenotypic Measurements in High-Content Imaging Screens. 161422 Preprint at 10.1101/161422 (2017).

22. Moon, K. R. et al. Visualizing Structure and Transitions in High-Dimensional Biological Data. Nature Biotechnology 37, 1482–92 (2019).

23. Traag, V. A., Waltman, L. & Eck, N. J. From Louvain to Leiden: Guaranteeing Well-Connected Communities. Scientific Reports 9, 5233 (2019).

24. Szklarczyk, D. et al. The STRING database in 2023: protein–protein association networks and functional enrichment analyses for any sequenced genome of interest. Nucleic Acids Res 51, D638–D646 (2023).

25. Steinkamp, R. et al. CORUM in 2024: Protein Complexes as Drug Targets. Nucleic Acids Research 53, 651–57 (2025).

26. Kanehisa, M., Furumichi, M., Sato, Y., Matsuura, Y. & Ishiguro-Watanabe, M. KEGG: Biological Systems Database as a Model of the Real World. Nucleic Acids Research 53, 672–77 (2025).

27. Hu, M. et al. Evaluation of large language models for discovery of gene set function. Nat Methods 22, 82–91 (2025).

28. Replogle, J. M. et al. Mapping information-rich genotype-phenotype landscapes with genome-scale Perturb-seq. Cell 185, 2559–2575.e28 (2022).

29. Rath, S. et al. MitoCarta3.0: an updated mitochondrial proteome now with sub-organelle localization and pathway annotations. Nucleic Acids Res 49, D1541–D1547 (2021).

30. Dixit, A. et al. Perturb-Seq: Dissecting Molecular Circuits with Scalable Single-Cell RNA Profiling of Pooled Genetic Screens. Cell 167, 1853–1866.e17 (2016).

31. Wainberg, M. et al. A genome-wide atlas of co-essential modules assigns function to uncharacterized genes. Nat Genet 53, 638–649 (2021).

32. Schaffer, L. V. et al. Multimodal cell maps as a foundation for structural and functional genomics. Nature 1–10 (2025) doi:10.1038/s41586-025-08878-3.

